# Transcriptional networks underpinning ploidy related increased leaf potassium in neo-tetraploids

**DOI:** 10.1101/2021.09.09.459632

**Authors:** Sina Fischer, Paulina Flis, Fang-Jie Zhao, David E. Salt

## Abstract

Neo-tetraploid *Arabidopsis thaliana* have elevated leaf potassium (K) driven by processes within the root. The root transcriptome of neo-tetraploids is distinct from diploids, with evidence of altered K homeostasis. Mutational analysis revealed that the canonical K-uptake transporters AKT1 and HAK5 are not required for this elevated leaf K in neo-tetraploids, while the endodermis, root hairs, and SOS signaling are. Contrasting the root transcriptomes of neo-tetraploids and diploids of mutants that block the neo-tetraploid K phenotype, allowed us to identify 91 differentially expressed genes associated with elevated leaf K in neo-tetraploids. This set of genes connects WGD to elevated leaf K, and is enriched in functions such as cell wall and Casparian strip development, and ion-transport, in the endodermis, root hairs, and procambium. This gene set provides tools to test the intriguing idea of recreating the physiological effects of WGD within a diploid genome.

## Main Text

It is well known that WGD events occurred multiple times throughout land plant evolution, suggesting an evolutionary benefit^1^. However, newly generated polyploids face severe problems, such as maintaining faithful chromosome separation during meiosis, or changes in cell architecture resulting from increased chromatin content^2^. The short term survival of neo-tetraploids was therefore speculated to occur at times of environmental change which would adversely affect diploid progenitors and neutralize their evolutionary advantage of being locally adapted^3^. Neo-tetraploid *A. thaliana*, rice (*Oryza sativa*) and citrange (*Citrus sinensis* L. Osb. x *Poncirus trifoliata* L. Raf.) all have increased tolerance to salinity and drought as a result of WGD, suggesting possible short term survival benefits of WGD^4–7^. Leaf K concentrations are higher in neo-tetraploid *A. thaliana*, driven by processes in the root as determined by grafting^4^, suggesting one possible mechanism of salinity tolerance through an enhanced ability to maintain K^+^/Na^+^ homeostasis under salinity stress^8^. Similarly, tetraploid rootstock-grafted watermelon (*Citrullus lanatus*) plants are more tolerant to salt stress than are diploid plants, and tetraploids also maintain better K^+^/Na^+^ homeostasis^9^. Neo-tetraploid *A. thaliana* and rice also show enhanced ABA and JA signaling, respectively, consistent with constitutive abiotic stress tolerance^5,10^. These observations suggest that under adverse conditions the negative impacts of WGD can be overcome, to provide a positive fitness benefit. However, although improved responses to abiotic stresses in polyploids have been documented, the exact molecular processes underlying these responses still remain to be discovered^11^.

To help address this gap and identify the molecular mechanisms linking WGD to improved stress tolerance we focused on the elevated leaf K that occurs in neo-tetraploid *A. thaliana*. Given that it has previously been established through micro-grafting that elevated leaf K of neo-tetraploids is driven by the root, we investigated the transcriptomic differences between roots from diploid and neo-tetraploid *A. thaliana* plants. We coupled this to the WGD of selected mutants with defects in genes necessary for elevated leaf K in neo-tetraploids. This allowed us to establish a causal link between changes in gene expression in roots of neo-tetraploids and elevated leaf K. Using this approach, we define a set of genes that provide functional insight into the underlying molecular mechanisms linking WGD and elevated leaf K. This gene set provides the tools needed to test the hypothesis that it is possible to recreate the physiological effects of WGD within a diploid genome. Leading the way to improved stress tolerance via simulating tetraploidy within a diploid genome.

## Results and Discussion

### Altered potassium homeostasis in neo-tetraploids

As previously reported, high leaf K concentration in neo-tetraploids is dependent on the ploidy status of the root^4^. To understand how neo-tetraploids modify K accumulation, gene expression in roots of diploids and neo-tetraploids was assessed. Plants were grown on agar solidified media under control conditions and with elevated NaCl. In line with previous results^4^, we observed elevated leaf K concentration in neo-tetraploids, and also the K chemical analogue rubidium (Rb) (Supplementary Fig. 1). Salinity stress intensifies the difference in leaf K concentration between diploid and neo-tetraploid plants (Supplementary Fig. 1), and revealed that neo-tetraploids have enhanced exclusion of Na from leaves, as previously observed in rice^4^.

We hypothesized that K transporter genes would be differentially expressed between diploids and neo-tetraploid roots. We selected a set of genes representative of known K transporters (genes identified at Thalemine https://bar.utoronto.ca/thalemine/begin.do, top hits) and performed a hierarchical clustering of our transcriptome data to show the expression pattern of this set of known K transporter genes in roots of both diploids and neo-tetraploids (Fig. 1a). The impact of salinity stress had overall the largest effect on the expression of the K transport genes, but within each treatment, gene expression was clearly differentiated between diploids and neo-tetraploids. Likewise, a PCA analysis using the K transporter genes showed that 64% of the variation in gene expression can be explained by the two factors of salinity stress (37%) and ploidy (27%) (Fig. 1b).

**Fig. 1:**
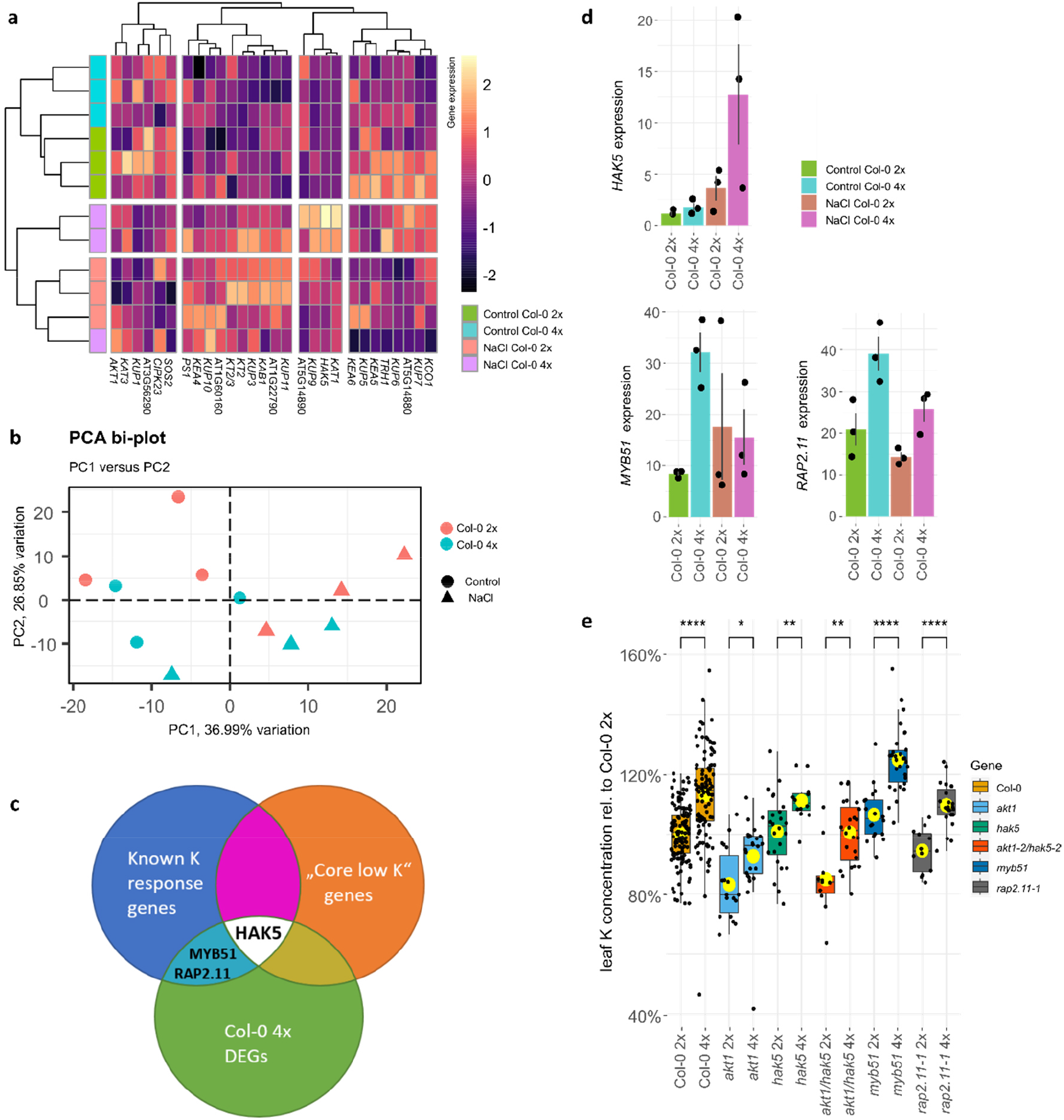
K genes in neo-tetraploid *A. thaliana*. **a)** RNAseq derived Gene expression pattern of 28 of 31 potassium genes taken from Thalemine (using the keyword search “K transporter” and restricting it to the 31 with highest scores) was measured in roots of diploid and neo-tetraploid wild-type. Plants were grown on ¼ Hoagland’s medium containing sucrose and agar with or without 35mM NaCl. A cluster analysis of samples from roots of diploid and tetraploid wild type plants shows a grouping based on ploidy. Genes with higher expression are shown with brighter colours. The annotation column identifies treatment and genotype. **b)** A PCA analysis of the same set of genes shows PC1 correlated with treatment and PC2 correlated with ploidy. Different colours indicate differences in ploidy and different symbols distinguish different treatments. Together they explained the majority (>63.84%) of the variation in expression of the 31 K-genes. **c)** The Venn diagram shows the intersections between DEG after WGD, low K treatment and known *HAK5* regulators. RNAseq analysis showed the enhanced expression of *HAK5* in neo-tetraploids which is also among the “Core K deficiency” response genes. In orange: “Core-low K”: 20 genes defined by meta analysis of expression studies^14–16^; In blue: 14 genes, known components of the low K signalling^12,13^; In green: 211 genes differentially expressed in wild type neo-tetraploid roots; **d)** Bar plots show *HAK5, MYB51 and RAP2*.*11* expression as analyzed by RNAseq (n=3, individual samples) in roots of diploid and neo-tetraploid wild type plants. DEGs were defined as having a fold change of >2 and a diverge probability of >0.8. **e)** Boxplot shows leaf K concentration in diploids and their neo-tetraploid counterparts normalized to diploid wild type. n=11-121 individual samples, data from 2 independent experiments using peat based soil in either jiffies^®^ or larger pots. Pairwise comparison using t-test indicates significant differences between diploids and neo-tetraploids. p-value ≤: *0.05, **0.01, ***0.001, ****0.0001 Yellow dot: averages Diploid (2x), neo-tetraploid (4x)

Having established that WGD affects the expression of K transporter genes in roots, we then tested if this difference observed in gene expression resembled a response to low external K. We compared all differentially expressed genes (DEGs) between diploids and neo-tetraploids with K response genes previously described^12,13^. Additionally, we defined a set of “Core low K” genes that have been found to be responsive to low K in the nutrient media in previously published studies^14–16^, and we also compared our DEGs to this set of “Core low K” genes. Only one gene, *High Affinity K Transporter 5* (*HAK5*) was found in all three sets (Fig. 1c,d; Supplementary Fig. 2). Additionally, two known *HAK5* regulators *MYB domain protein 51* (*MYB51*) and *Related to AP2 11* (*RAP2*.*11*)^12,13^ are DEGs in neo-tetraploids (Fig. 1d). We conclude that although WGD causes a clear K response in roots of neo-tetraploids, this response does not resemble a typical transcriptional response to low supply of K in the nutrient media. We speculate that instead of a response to low K supply, which is what has previously been studied^17–20^, the different K response in neo-tetraploid roots may be driven by increased internal demand for K, required to meet the needs of the larger cell size of neo-tetraploids ^21,22^ (Supplementary Fig. 3).

To assess the impact of *HAK5* and its regulators *MYB51* and *RAP2*.*11* on K concentration in leaves of neo-tetraploids, knock out mutants were obtained, and their genomes doubled. *A. thaliana* has two major K uptake systems, the high affinity system which is mediated by *HAK5* and the low affinity system for which *K*^*+*^ *Transporter 1* (*AKT1*) is a major component^23,24^. We assessed the effect of a lack of both systems on the leaf K concentration in neo-tetraploids grown in soil by investigating *akt1-2* and *hak5* mutants. To account for redundancy between these two transporters we also generated the neo-tetraploid of the *akt1-2 hak5-2* double mutant^25^. We assessed the leaf ionome in these genotypes in each case for diploids and neo-tetraploids. A two-way ANOVA of the leaf K concentrations showed that genotype (i.e. *hak5, akt1-2*, wild type etc) has a significant impact (p value <2e-16), as did ploidy (i.e. 2x and 4x, p value <2e-16). However, there was no significant interaction between ploidy and genotype (p value 0.4248). We performed a t-test to directly compare lines before and after WGD. We observed that wild type neo-tetraploids show an increased leaf K concentration (Fig 2e), as expected^4^. However, loss of function of neither *AKT1* nor *HAK5* had an impact on the increased leaf K concentration in neo-tetraploids, with all neo-tetraploid lines showing significantly higher leaf K (Fig 2e) and Rb (Fig S4) concentrations than their diploid progenitors. We also observed smaller variations in the concentration of some other mineral nutrients (Fig. 1e & Supplementary Fig. 4). This supports the conclusion that elevated leaf K in neo-tetraploids is not simply due to enhanced activity of the known K-uptake systems of HAK5 and AKT1.

**Fig. 2:**
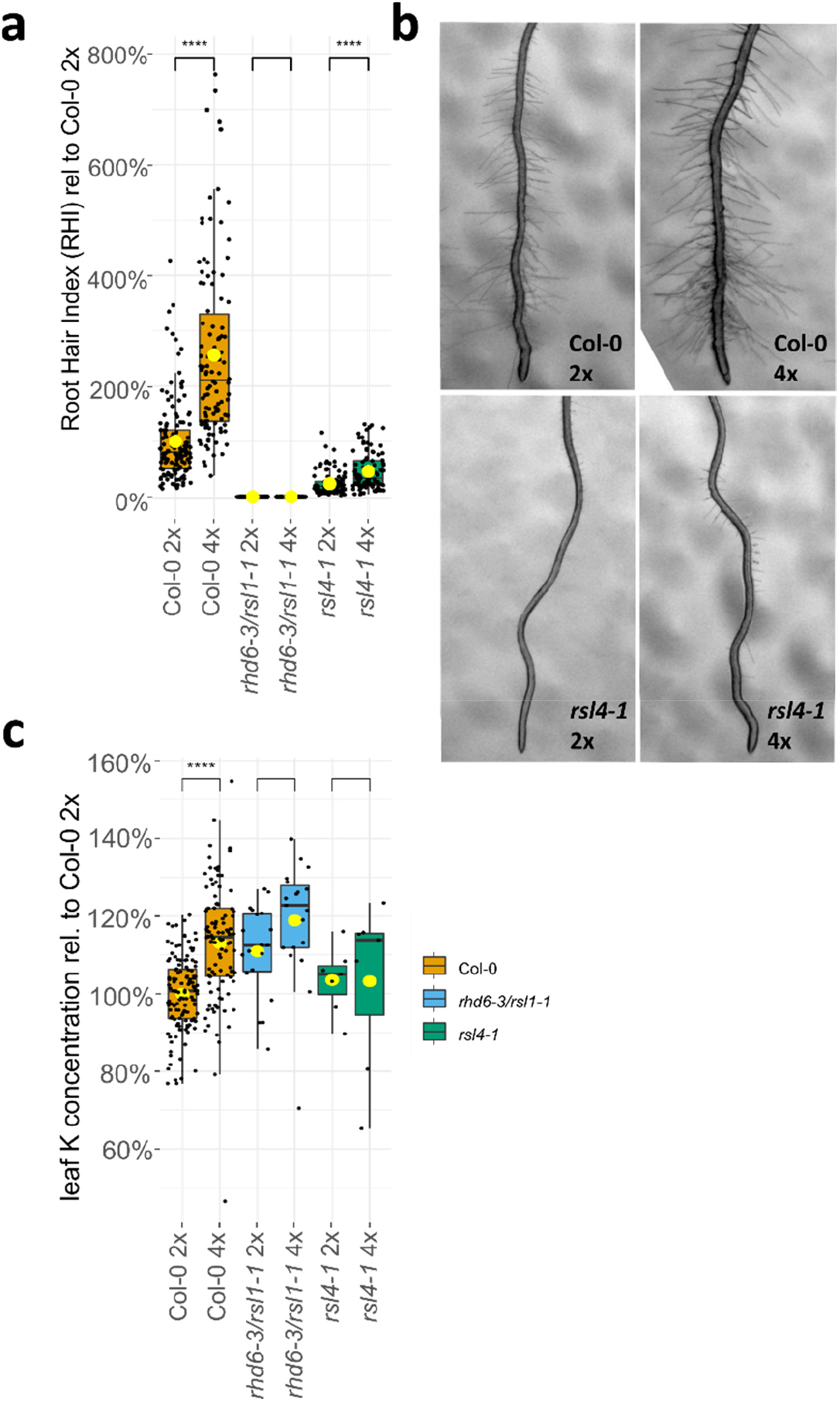
Root hairs of neo-tetraploids grown on 1/10 Hoagland without Sucrose. **a)** Boxplot of RHI relative to diploid wild type shows longer and denser root hairs in neo-tetraploid lines except for the *rhd6-2/rsl1-1* mutant which does not have any root hairs. n=51-134, individual samples. Plants were grown on 1/10 Hoagland’s medium with high purity agarose and without sucrose. **b)** Microscope images showing root hairs. Measurements were taken 1cm above the root tip. Root hair density and length was assessed. A product of root hair density and length constitutes the RHI. **c)** Relative leaf K content of soil grown plants shows suppression of the leaf K phenotype in root hair mutants. n= 7-109, individual samples, Pairwise comparison using t-test indicates significant differences between diploids and neo-tetraploids. p-value ≤: *0.05, **0.01, ***0.001, ****0.0001 Yellow dot: averages. Diploid (2x), neo-tetraploid (4x)

### Impact of root hairs on leaf K in neo-tetraploids

Root hairs play an important role in K uptake in diploid *A. thaliana*^26^, and neo-tetraploid *A. thaliana* are known to have longer and denser root hairs^27^. We therefore directly tested the role of root hairs in the elevated leaf K of neo-tetraploids. First, we analyzed the transcriptional profiles of 106 “root hair” genes (Thalemine key word search) from our root transcriptome data using hierarchical clustering. We found expression patterns based on Na treatment, and within those, based on ploidy (Supplementary Fig. 5). Further, two “root hair” genes, *Leucin-rich Repeat/eXtensin 1* and *Root Hair Specific 13* are DEGs in neo-tetraploids. This evidence is consistent with a role of the root hairs regulatory network in controlling leaf K accumulation in neo-tetraploids.

Then, we assessed the root hair index (RHI), the product of multiplying the length of the root hairs by the density of the hairs in the root, in wild type and known root hair mutants *rsl4-1* and *rhd6-3/rsl1-1* before and after WGD. We found an increase in RHI in wild type neo-tetraploids (Fig. 2a,b) along with an increase in leaf K (Fig. 2c), as expected. In contrast, in the double mutant *rhd6-3/rsl1-1* we observe a complete lack of root hairs in the diploid^28^, and also after WGD (Fig. 2a). In the root hair mutant *rsl4-1* we observe a reduction in RHI in the diploid mutant *rsl4-1*^29^ but after WGD we observe an increase in root hairs in *rsl4-1* but to levels lower than observed in the wild type (Fig. 2a,b). Importantly, loss or reduction of RHI in *rhd6-3/rsl1-1* and *rsl4-1*, respectively, abolishes the increase in leaf K observed in wild type after WGD in plants grown in soil (Fig. 2c). A two-way ANOVA showed an impact of both ploidy (*p* value <2e-16) and genotype (*p* value <2e-16) on the RHI, as well as for their interaction (*p* value 2.25e-13). In contrast, the leaf K concentration of these lines showed an impact for ploidy (*p* value 2.01e-9) and genotype (*p* value 0.001028), but not their interaction (*p* value 0.11307). The pair wise comparison using a t-test reveals significant differences in leaf K only for the wild type (Fig. 2c). Both *rsl4-1* and *rhd6-3/rsl1-1* blocked any increase in leaf K in the neo-tetraploids. Further, we assessed the RHI of other root hair mutants such as *rhd1-2, rhd2-4, rsl1-1* and *rsl2-1*. We did not observe any decrease in RHI in diploids of these mutants compared to wild type (Supplementary Fig. 6a), and they had no impact on ploidy related leaf K (Supplementary Fig. 6b).

We therefore establish, using two independent mutants, that a reduction in root hairs results in a suppression of the neo-tetraploid leaf K phenotype. This indicates that while disrupting individual K uptake components such as HAK5, AKT1 or their regulators has no effect on the increased leaf K of neo-tetraploids, loss of a major K uptake cell-type, root hairs, does abolish this phenotype. This suggests significant redundancy in K-uptake mechanisms within root hairs beyond HAK5 and AKT1.

### A role for ABA signaling

Neo-tetraploid *A. thaliana* have positively regulated ABA signaling, suggesting they have a pre-activated ABA response, which is consistent with their enhanced stomatal closure, salinity and drought tolerance^5^. K homeostasis is interconnected with ABA signaling, guard cell movement and salinity tolerance^30^. Therefore, to identify causal genes involved in elevated leaf K in neo-tetraploids we utilized our RNAseq dataset of wild type diploids and neo-tetraploids to identify DEGs in roots involved in ABA signaling. A GO enrichment analysis revealed several GO terms overrepresented among the DEGs in wild type neo-tetraploids grown under control conditions, including response to ABA, consistent with what has previously been observed^5^ (Fig. 3a). These GO terms are only enriched when plants are grown in the absence of elevated salinity. This suggests that neo-tetraploids may be pre-adapted to abiotic stress as previously proposed^5^. Due to their potential link with K homeostasis we focus on the 11 DEGs within the ABA enrichment group (Fig 3).

**Fig. 3:**
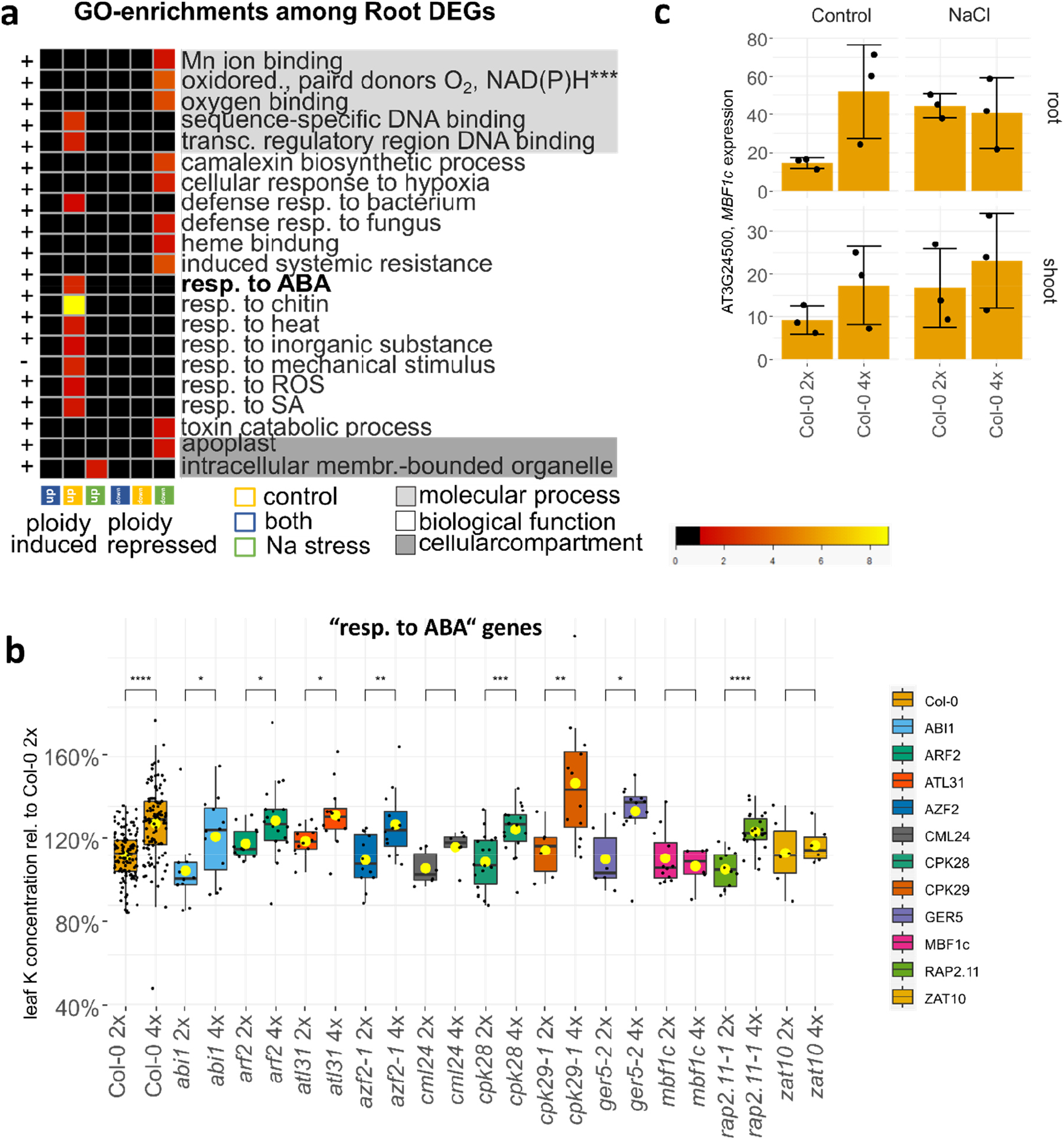
Role of ABA-signaling. **a)** Heatmap shows the p-value of significantly enriched GO terms among genes differentially expressed (DEGs) after WGD in plants grown for two weeks on ¼ Hoagland’s, agar solidified media containing sucrose and 0 or 35mM NaCl. For gene selection see Supplementary Fig. 11. Among the enriched classes is “response to ABA” containing 11 genes. It is overrepresented among genes induced in neo-tetraploid roots and is connected to tolerance to abiotic stress. Analysis was done using PANTHER. Displayed are only the most defined terms or child terms. A full list can be found in Table S2. Test Type: Fisher’s Exact, Bonferroni correction for multiple testing, GO database release 2019-12-09. Abbreviations for GO categories: ***=oxidoreductase activity, acting on paired donors, with incorporation or reduction of molecular oxygen, NAD(P)H as one donor, and incorporation of one atom of oxygen, transc.=transcription, resp.=response, membr.=membrane **b)** Boxplot shows leaf K concentration in diploids and their neo-tetraploid counterparts. Data is normalized to the diploid wild type. Plants were grown in soil for 5 weeks. n=6-102, individual samples, data from 2 independent experiments using peat based soil in either jiffies^®^ or larger pots. Two way ANOVA shows significant effects of ploidy and ploidy*genotype[MBF1c] on the leaf K concentration. Pairwise comparison using t-test indicates significant differences between parental lines and crosses. p-value ≤: *0.05, **0.01, ***0.001, ****0.0001 Yellow dot: averages. **c)** Bar plots show the RNAseq assessed expression (mean Fragments Per Kilobase of transcript per Million mapped reads (FPKM)±SD) of *MBF1c* in plants grown on agar solidified ¼ Hoagland with sucrose, with and without salinity stress (35mM NaCl) in root and shoot tissues. n=3, individual samples. Significant increase in expression in roots of control grown plants (cutoff fold change ≥2 and a diverge probability ≥0.8). Diploid (2x), neo-tetraploid (4x)

We were able to isolate homozygous T-DNA insertion alleles for 10 of these 11 genes, and generated neo-tetraploids. We also included the ABA receptor *ABI1*. We then assessed the leaf ionome of these neo-tetraploids grown in soil. A two-way ANOVA of the leaf K concentration (Fig. 3b) of these plants showed a significant interaction only for ploidy and the *Multiprotein Bridging Factor 1c* (*MBF1c*) gene. The *mbf1c* diploid and neo-tetraploid mutant showed no difference in their leaf K concentration. Further, *cml24* and *zat10* also showed no significant difference in leaf K between diploid and neo-tetraploid, though in *cml24* the trend is visible. *MBF1c* is one of three MBF genes in *A. thaliana*. MBF’s are transcriptional co-activators^31^. They are highly conserved among eukaryotes^32^. In *A. thaliana MBF1a* and *1b* show high sequence similarities and expression profiles^33^. They are thought to be important for pathogen response. *MBF1c* however is distinct from *a* and *b*. It is induced by ABA treatment and has been shown to be involved in mediating salinity and heat stress^32,34^. In neo-tetraploids *MBF1c* shows higher expression in roots only in plants grown under control conditions (Fig. 3c). Upon Na treatment *MBF1c* is induced in roots of wild type diploid plants. However, in neo-tetraploids *MBF1c* is constitutively activated to the levels seen in the Na-treated diploids, with Na treatment having no impact on expression in neo-tetraploids. Identification of genes induced in an *MBF1c* ectopic over expression line revealed potential targets for co-regulation through MBF1c^34^. Among these 87 genes, 18 are DEGs in roots of wild type neo-tetraploids when grown under control conditions, supporting a role for MBF1c in the transcriptional response of roots to WGD. MBF1c sits upstream of several ABA and ethylene response genes, and is essential for the establishment of the elevated leaf K observed in wild type neo-tetraploids. *ZAT10* is a transcriptional repressor involved in abiotic stress responses. Both its constitutive expression and knockout has been shown to result in increased osmotic and salinity stress, suggesting it modulates abiotic stress responses^35^. However, other ABA response genes known to be directly involved in ABA signaling such as *ABI1* whose mutant *abi1-1* is insensitive to ABA^36^, are not required for elevated leaf K in neo-tetraploids as loss of function mutants in these genes still show elevated leaf K in neo-tetraploids (Fig 3b). This suggests that there is not a direct link between ABA signaling and elevated leaf K in neo-tetraploids.

### Other candidate genes potentially involved in the regulation of the elevated leaf K concentration in neo-tetraploids

To further explore other possible genes involved in the neo-tetraploid K phenotype we selected a further 20 additional genes from the literature that had the potential to impact leaf K concentration of neo-tetraploids. For example, *High-Affinity K+ Transporter1* (*HKT1*), a Na-transporter that has been shown to be important for K^+^/Na^+^ homeostasis during salinity stress^8^. These selected candidate genes were grouped, based on their function, into seven groups (Fig. 4).

**Fig. 4:**
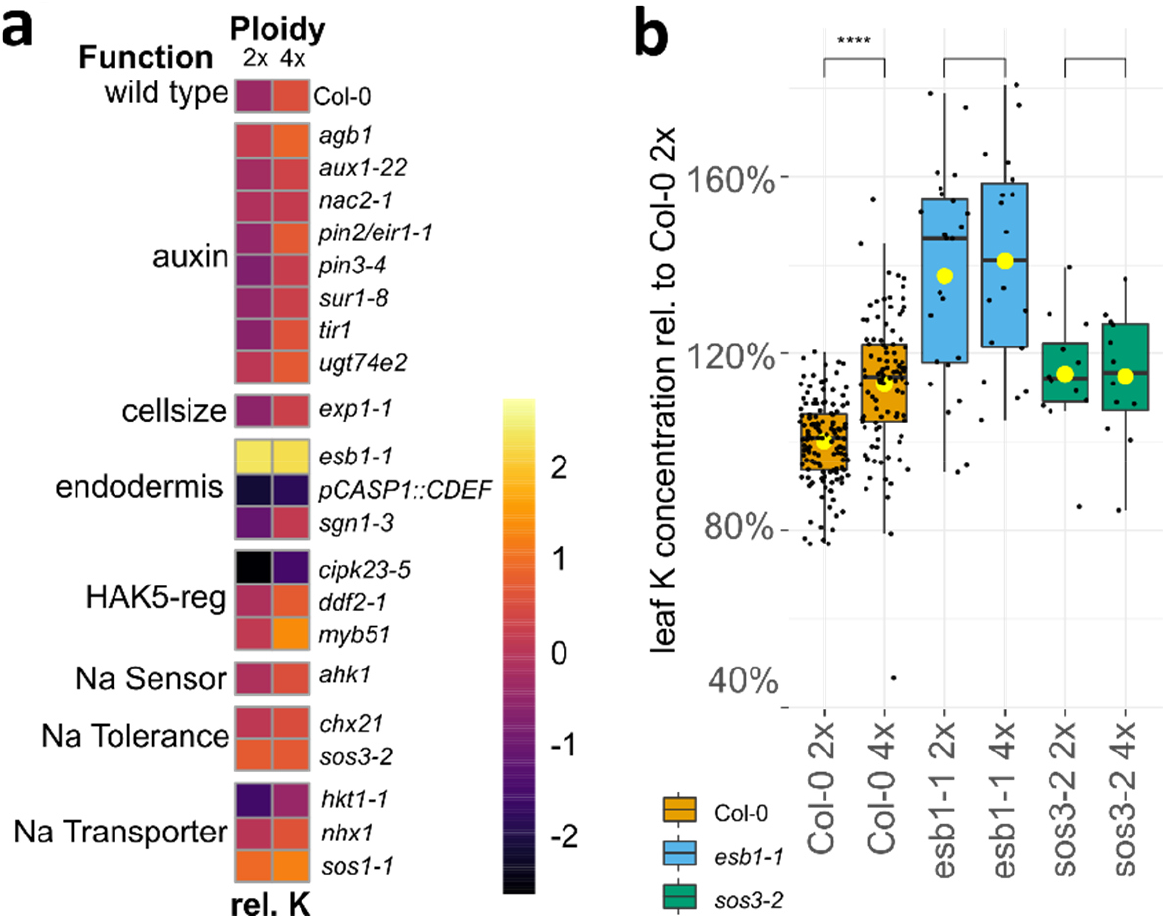
Selected candidates genes. **a)** Heatmap displays K concentration in leaves of diploid and neo-tetraploid mutants grown on soil relative to the K concentration in diploid wild type plants. The color scale shows genotypes with lower (darker) and higher (brighter) than average relative K concentrations within each functional group annotated using previously published studies. The annotation rows show their function and ploidy. **b)** Boxplot shows the leaf K concentration relative to diploid wild type of the 2 mutants, *esb1-1* and *sos3-2* for which a significant interaction between ploidy and gene was found in the 2 way ANOVA. n= 6-109, individual samples, Pairwise comparison using t-test indicates significant differences between diploids and neo-tetraploids. p-value ≤: *0.05, **0.01, ***0.001, ****0.0001 Yellow dot: averages. Diploid (2x), neo-tetraploid (4x)

We generated neo-tetraploids for loss-of-function mutants for all these genes and other lines, and assessed the leaf ionomes of soil grown plants. Fig. 4a shows the leaf K concentration relative to that of the diploid wild type for all these mutants and lines, grouped by function. For most functional groups, an increase in leaf K from diploids to neo-tetraploids is apparent, showing a typical pattern of low (purple) to high (orange) leaf K, for example the “auxin” group. However, for both the “endodermis” and the “Na tolerance” functional groups this pattern is disturbed. In the “endodermis” functional group, we see high and low leaf K concentrations among both diploid and neo-tetraploids. In the “Na tolerance” functional group on the other hand leaf K concentrations are generally high with no color change, signaling no increased K in the neo-tetraploids. A two-way ANOVA shows significant changes in leaf K for ploidy (p value <2e-16) and gene (p value <2e-16) but not their interaction (p value 0.3563). A pairwise comparison indicates a significant effect only for ploidy[4x] * gene[*ESB1*] (p value = 0.007) and for ploidy[4x] * gene[*SOS3*] (p value = 0.024). For both *esb1-1* and *sos3-2* the increased leaf K concentration normally observed in the neo-tetraploids is suppressed (Fig. 4b). A two-way ANOVA showed both factors, ploidy and gene, as well as their interaction now to be significant in affecting the leaf K concentration in wild type, *esb1-1* and *sos3-2* (p values 1.48e-6, <2.2e-16 and 0.0076 respectively). We assessed if this loss of elevated leaf K in *esb1-1* and *sos3-2* neo-tetraploids could be due to a reduction of root hairs, as seen for *rsl4-1* and *rhd6-3/rsl1-1*. However, both *esb1-1* and *sos3-2* neo-tetraploids exhibit a higher RHI than their diploid progenitor (Supplementary Fig. 6c), establishing that suppression of elevated leaf K in neo-tetraploids of *esb1-1* and *sos3-2* is not related to changes in root hairs.

*SOS3* (also known as *CBL4*) encodes a Ca sensor which operates in the SOS signaling network in response to salinity stress^37^. SOS3 facilitates Na^+^ efflux, leading to reduced K^+^ efflux through depolarization-activated K Outward Rectifying channels (KOR)^38^. WGD may require SOS signaling to enable enhanced K accumulation even on low external Na concentrations. *ESB1* on the other hand is an important component for the biosynthesis of Casparian strips, which form in the cell wall at the endodermis^39^. In *esb1-1* Casparian strips are disturbed, leading to an increase in leaf K concentrations. It is thought this increase in leaf K is due to the enhanced endodermal suberization observed in this mutant^39^. Increasing the suberized zone of the endodermis prevents K leaking from the stele where it is more highly concentrated than in the surrounding root tissue^40^. WGD of *esb1-1* does not lead to a further increase of leaf K in the neo-tetraploids. This suggests that a normally functioning endodermis plays an important role in the enhanced leaf K concentration we observe in wild type neo-tetraploids.

### Transcriptome of mutants in genes necessary for elevated leaf K in neo-tetraploids

To further probe the gene expression changes required for increased leaf K in neo-tetraploids we utilized the three mutants *sos3-2, esb1-1* and *rhd6-3/rsl1-1* we had established to suppress elevated leaf K in neo-tetraploids. We grew diploid and neo-tetraploid mutants and wild type on soil, confirmed their leaf K phenotypes (Supplementary Fig. 7), and performed a transcriptomic analysis. We identified DEGs in roots between neo-tetraploids and their diploid progenitors. Wild type had 114 DEGs, and this number was reduced in all three mutants suppressing the K ploidy phenotype (Fig. 5a). We next compared all DEGs between diploids and neo-tetraploids across the wild type and three K phenotype suppression mutants (Fig. 5b). Through this comparison we identified 91 genes specifically responsive to WGD in wild type that are associated with the elevated leaf K concentration after WGD. A heat map of the expression of these 91 genes in wild type identified two clusters, one contains 17 genes that are up regulated, and the other 74 genes that are down regulated by WGD (Fig. 5c). A GO enrichment analysis revealed enrichment of genes involved in xyloglucan processes (cell wall), ion transport (zinc and inorganic anions) and the Casparian strip (Fig. 5d).

**Fig. 5:**
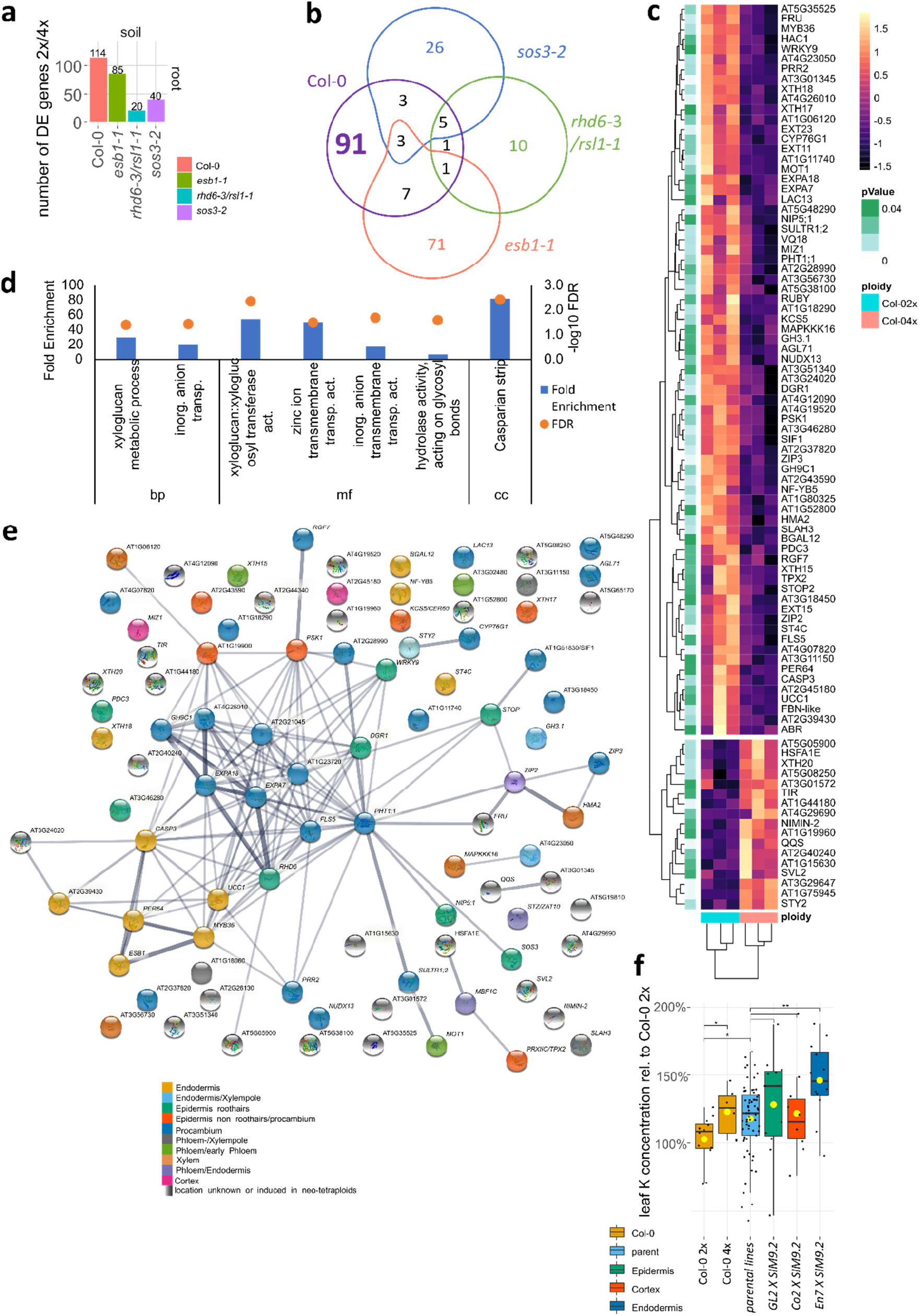
Neo-tetraploid elevated leaf K gene network. **a)** Number of ploidy related gene expression changes (DEGs) in roots of soil grown plants shows fewer differences in neo-tetraploid mutants supressing the elevated leaf K phenotype of neo-tetraploids. **b)** Venn comparison of all ploidy dependent DEGs reveals that 91 are specifically changed in wild type but not in the K-phenotype supressing mutants. **c)** Heatmap of the expression of 91 genes in diploid and neo-tetraploid wild type shows a clear ploidy pattern with 17 genes induced in neo-tetraploids and 74 genes repressed. The annotation row indicates significance of fold change of expression between diploid and neo-tetraploid wild type. Dark: less significant, light: highly significant **d)** Graph displaying fold enrichment and significance (-log10 false discovery rate FDR) of GO categories among the 91 genes. Analysis was done using PANTHER. Displayed are only the most defined terms or child terms. A full list can be found in Table S2. Test Type: Fisher’s Exact, Bonferroni correction for multiple testing, GO database release 2019-12-09. **e)** STING network analysis of 74 down regulated genes plus 4 previously identified genes known to be related to the elevated leaf K phenotype of neo-tetraploids. The STRING network identified hub genes, connected through co-expression correlations with several other genes of the ploidy network. Colours are chosen based on root cell type specific expression as shown in the public database ePlant. A heatmap for the colour annotation can be found in Supplementary Fig. 8. For genes without colour no data was available on ePlant. **f)** Boxplot shows the leaf K concentration relative to diploid wild type of lines with increase ploidy in epidermis (GL2), cortex (Co2) or endodermis (En7) only ^54^. Ploidy increase was achieved by expression of the endoreduplication gene SIM under control of a tissue specific promotor. Crosses containing both the promotor and SIM construct display increased endoreduplication i.e. polyploidy in one of the 3 cell types. n=6-65, individual samples. Pairwise comparison using t-test indicates significant differences between parental lines and crosses. p-value ≤: *0.05, **0.01, ***0.001, ****0.0001 Yellow dot: averages. Diploid (2x), neo-tetraploid (4x)

To further understand the function of these genes in the K ploidy phenotype we investigated their cell-type specific expression patterns. We utilized a publicly available database (ePlant) to search for the cell type specific expression pattern of all 74 genes repressed by WGD in wild type neo-tetraploid roots. We focused on genes which are repressed because we reasoned that this repression must have occurred in the cell type the gene is usually expressed in. For genes which are induced we cannot make any assumptions about where they may be induced in neo-tetraploids, and therefore they were excluded from this analysis. We also included the genes we have already established as being necessary for elevated leaf K in neo-tetraploids, which are *MBF1c, ZAT10, SOS3, ESB1, RHD6* and *RSL1*. Root cell type specific data was available for 68 of the 80 genes. We generated a heat map based on their tissue specific expression pattern and clustered them into 10 groups. Each cluster defined a pattern of cell-type specific gene expression within the root (Supplementary Fig. 8). We detected a cluster of 9 genes, repressed in neo-tetraploids, which are mainly expressed in the endodermis. In this cluster we find *ESB1* which we had already established as being necessary for the ploidy K phenotype. This cluster contains several genes known to be involved in Casparian strip formation, including *MYB36* the master transcriptional regulator, and *PER64, UCC1* and *CASP3* regulated by *MYB36*^41–44^. Very closely clustered are two more groups of 2 genes each. In one, genes are additionally expressed in the xylem pole. For these, a direct involvement in Casparian strip formation has not been shown. It includes *GH3*.*1* which is involved in auxin signaling^45^. In the other, the two genes *MBF1c* and *ZAT10*, which we had confirmed to be important for the K phenotype (Fig 3b), are additionally expressed in the phloem. Two further clusters encompass genes expressed in the epidermis, some of them in root hair cells, and others in non-root hair cells. Notable here is the *WRKY9* transcription factor which has recently been shown to be induced by salt treatment, and to positively regulate *CYP94B3* and *CYP86B1*, leading to increased root suberin and salt tolerance^46^. A large cluster of genes is expressed in the procambium, among them *LAC13* which has recently been shown to also localize to the Casparian strip region^47^. Several genes in the procambium cluster are also expressed in the epidermis. For example, *EXPA7/Expα-1*.*26*, codes for an expansin. Expansins are a group of proteins involved in cell wall loosening and turgor driven cell wall expansion^48^. Expression levels of *EXPA7* are positively correlated with root hair length^49^. The last clusters encompass stele and cortex specific genes. Changes in expression in *MAPK*^*3*^*16*, which is down regulated in neo-tetraploids, may alter a signaling pathway which could in turn affect K translocation to the shoot. MAPK^3^16 is expressed in the xylem (Supplementary Fig. 8). We used this clustering to assign colors to genes in a network analysis.

Using STRING^50^ we assembled a network of the 91 genes differentially expressed in wild type neo-tetraploids and associated with elevated leaf K, and also included the six genes we had established to affect the leaf K phenotype in neo-tetraploids (Fig. 5e). In the network genes are connected based on their co-expression characteristics. We detect several hub genes; genes co-expressed with numerous other genes. One of these hubs is *EXPA7*, expressed in the procambium and in root hair epidermal cells. *EXPA7* is connected to several endodermal genes, and other root hair and non-root hair genes. Among them is *PHT1;1*, which encodes for one of two major phosphate uptake transporter in roots, the other one being *PHT1;4*. Both play a role in Pi acquisition from low and high Pi environments and a lack of both genes leads to increased root hair length^51^. *PHT1;1* is very highly expressed in root hair cells^52^ which are important for Pi uptake. This shows that the most prominent cluster is centered around root hairs and cell growth, again showing the importance of root hairs for the establishment of the K phenotype in neo-tetraploids. Another hub gene is *MYB36* which is connected to, as expected, many Casparian strip genes, but also several genes expressed in the procambium. This network (Fig. 5e) represents gene expression changes in roots of neo-tetraploids induced by WGD and associated with elevated leaf K. A subset of these genes will be necessary for the K ploidy phenotype, while other are likely to be pleotropic side effects induced by expression changes in these causal genes. For example, root hairs are necessary for the K ploidy phenotype (Fig. 2), but not all genes involved in root hair function will be causal for the phenotype. It is unlikely for example that the *PHT1;1* hub is required for the K ploidy phenotype; its expression is changing in neo-tetraploids because root hairs are changing.

### Root cell-type specificity of WGD drive increased leaf K

In order to further narrow down the set of genes described in the co-expression network (Fig. 5e), we asked the question, what cell-types in the root need to undergo WGD to initiate the K ploidy phenotype? To answer this, we utilized the ectopic expression of *SIM* in a tissue specific manner. *SIM* is sufficient for endoreduplication^53^, the replication of the nuclear genome in the absence of mitosis, which leads to elevated nuclear gene content and polyploidy. Expression of *SIM* in a cell-type specific manner allows targeted endoreduplication in specific cell-types^54^. Constitutive overexpression of *SIM* is very disruptive to growth. Previously a system using *GAL4-VP16* driven transactivation was established to assess the effect of tissue specific overexpression in a heterozygous F1 generation^54^. Tissue specific promotor lines *pEn7-GAL4-pUAS-H2HF, pGo2-GAL4-pUAS-H2AF, pGL2-GAL4-pUAS-H2AF*, and *pUAS* controlled *SIM* lines *pUAS-SIM3*.*3, pUAS-SIM9*.*2* were crossed to generate a tissue specific increase in ploidy. Nuclear localized GFP was also expressed under control of the tissue specific promotors, illustrating the location of SIM expression in the root. Using this approach, we were able to drive endoreduplication specifically in the endodermis (promoter *pEn7*), cortex (promoter *pCo2*) and epidermis (promoter *pGL2*). An analysis of K in leaves of these lines grown in soil revealed that endoreduplication in the endodermis (line *pEn7 X SIM9*.*2*) produced increased leaf K concentration compared to the parental lines (Fig. 5f). However, we observed no increase in leaf K when endoreduplication is targeted at the cortex (lines *pGL2 x SIM9*.*2*) or epidermis (lines *pCo2 x SIM9*.*2*) (Fig. 5f). Further, crosses of the cell-type specific promoter lines with two other independent *SIM* lines *SIM1*.*1* and *SIM3*.*3* did not show effects on leaf K (Supplementary Fig. 10a). This can be attributed to the silencing to the *SIM* transgene in *SIM1*.*1* which shows no SIM related changes in phenotype (Supplementary Fig. 9), nuclei or cell size. *SIM3*.*3* is likely to have weak SIM expression as reflected in the weaker GFP signal (Supplementary Fig. 10b-d) which could explain why *pEn7 X SIM9*.*2* has a stronger impact in leaf K than *pEn7 X SIM3*.*3*.

We examined the root hairs of these *SIM* lines to assess if an increase in root hairs is responsible for the K phenotypes of *En7 X SIM9*.*2* with endoreduplication specifically in the endodermis. *En7 X SIM9*.*2* does have somewhat earlier root hair formation, though this slight increase in root hairs is not as strong as in other SIM lines that do not displaying elevated leaf K (Supplementary Fig. 10e). We conclude that the elevated leaf K observed in the *En7 X SIM9*.*2* line with endoreduplication of the endodermis is not due to increased root hairs. Root hairs are necessary for increased leaf K in neo-tetraploids but they are not sufficient. Further, we conclude that WGD in the endodermis is sufficient to produce the elevated leaf K observed in wild type neo-tetraploids.

## Summary

We discovered that disruption of a singular K-uptake gene, or even the two major high and low affinity K uptake systems or their regulators, is not enough to suppress the elevated leaf K phenotype of neo-tetraploids. This suggests significant undiscovered redundancy within the K uptake system. We show genetically that roots hairs (a main site of K uptake), a functional endodermis, SOS signaling, and certain aspects of ABA signaling are necessary for the ploidy K phenotype. Expanding our approach beyond single genes, we uncover a transcriptional network associated with the increased leaf K concentration in neo-tetraploids. This network contains genes that are both repressed or activated in neo-tetraploid roots, and is enriched in processes such as root hair development, cell wall re-modeling, ion-transport, endodermal development and ABA signaling. These genes are differentially expressed in key cell-types of the root, including procambium, epidermis, root hairs, endodermis and vascular tissue. WGD appears to initiate root-wide changes in gene expression that lead to elevated leaf K. However, we observe that WGD within the endodermis alone is sufficient to drive elevated leaf K concentrations. It remains to be determined which genes within this root-wide transcriptional network are necessary for elevated leaf K in neo-tetraploids, and which are sufficient. Further, it is an open question as to what the molecular steps initiated by WGD in the endodermis are that lead to root-wide transcriptional changes driving elevated leaf K in neo-tetraploids. The gene set we have identified should help answer these questions. In certain crops^10,55,56^ tetraploids are more tolerant to drought and salinity. This gene set offers the potential to synthetically recreate, in a diploid background, these beneficial effects of WGD without the associated drawbacks, providing a new avenue to develop crops more resistant to drought and salinity.

## Materials and Methods

Lines were obtained from NASC, or donated. A full list of can be found in Table S1. Elemental content was determined using ICP-MS, image analysis was done using Fiji and statistical analysis and graphs were done using R/RStudio. A full description of the material and methods can be found in the supplemental materials. RNAseq was performed by BGI (https://www.bgi.com/global/) according to their standard analysis procedure and by Deep Seq at the University of Nottingham (https://www.nottingham.ac.uk/deepseq/).

## Supporting information

Supplemental Table 1 - Plant Lines

Supplemental Table 2 - GO Enrichment

## Availability Statement

RNAseq datasets generated during this study can be found at GEO https://www.ncbi.nlm.nih.gov/geo/ under the number GSE180004 for the assessment of plate grown wild type plants and GSE180818 for the assessment of soil grown wild type and mutant plants. The latter dataset contains further samples not described in this study of plate grown plants and a fifth mutant chx23-4. Both sets have also been unified under the SuperSeries GSE180819.

## Acknowledgments

We thank Francisco Rubio for his generous donation of a set of *hak5-3, akt1-2, cipk23-5, akt1-2/cipk23-5, akt1-2/hak5-2, hak5-3/cipk23-5* and the *akt1-2/cipk23-5/hak5-3* triple mutant lines ^25^. The lines *rhd1-2, rhd2-4, rhd6-3/rsl1-1, rsl1-1* and *rsl4-1* were donated by Rahul Bhosale. The line *CASP1-GFP* by Guilhem Reyt whom we also thank for is help with confocal microscopy. We thank Daniela Dietrich for donating the *GL2, Co2, En7 X SIM* crosses^54^. We thank Priya Ramakrishna for the donation of *expa1-1* seeds. Lines *aux1-22* and *pin3-4* were donated by Malcolm Bennett.

This study was supported by funding to SF from the DFG (FI 2152/1-1, DFG Fellowship), the Royal Society (Research Grant, RGS\R1\201381) and the University of Nottingham (Nottingham Research Fellowship and the Future Food Beacon of Excellence). We acknowledge a grant to F-JZ and DES from National Science Foundation Council of China (NSFC) International Collaborative Project.

