## Supplemental Figures and Methods for "Transcriptional networks underpinning ploidy related increased leaf potassium in neo-tetraploids"

Sina Fischer1, Paulina Flis 1,Fang-Jie Zhao2,David E. Salt1*
1Future Food Beacon of Excellence and the School of Biosciences, University of Nottingham, Nottingham LE12 5RD, UK
2State Key Laboratory of Crop Genetics and Germplasm Enhancement, College of Resources and Environmental Sciences, Nanjing Agricultural University, Nanjing, China

*David E. Salt;

**This PDF file includes:**

Supplementary text: Materials and Methods (Extended)
Figures S1 to S14
SI References

**Supplementary Information Text**

**Materials and Methods (Extended).**

### *Arabidopsis thaliana* lines

*A. thaliana* lines were obtained from the Nottingham Arabidopsis Stock Center (NASC) and donated by Francisco Rubio[1], Daniela Dietrich[2], Guilhem Reyt[3], Priya Ramakrishna[4]. The wild type Col-0 tetraploid lines were obtained from Chao *et al*.[5]. Further, previously published lines were donated by Rahul Bhosale and Malcolm Bennett[6–10].

### Cultivation conditions

**Agar solidified nutrient media**: Plants were grown on ½ MS (2.2g MS/l, 1% agar, 5mM MES), ¼ or $^{1}/_{10}$ Hoagland medium (250/ 100 µM $NH_4H_2PO_4$, 500/ 200 µM $MgSO_4$, 700/ 280 µM $Ca(NO_3)_2$, 1500/ 600 µM $KNO_3$, 12.5/ 5 µM Fe-HBED, 5 mM MES at pH 5.7.) solidified with 1 % (w/v) agar (Type A, Sigma Aldrich or Alfa Aesar Low EEO Agarose for high purity agar). In some cases the nutrient solution contained and 1 %w/v sucrose (Sigma Aldrich BioXtra). Cultivation was under long day conditions of 16h light (170-230 µmol $m^{-2}$ $s^{-1}$) /8h dark.

**Soil experiments:** Plants cultivated on soil was carried out as described previously[11,12]. In brief, plants were grown for 5 weeks on peat Jiffy® soil pellets or in pots filled with peat substrate (Levington M3, T34 biocontrol®) after stratification for 48h at 4°C. Genotypes were randomized. Plants were bottom watered with nutrient solution (100 µM $NH_4H_2PO_4$, 200 µM $MgSO_4$, 400 µM

$Ca(NO_3)_2$, 600 µM $KNO_3$, 5 µM Fe-HBED, 4.63 µM $H_3BO_3$, 0.032 µM $CuSO_4$, 0.915 µM $MnCl_2$, 0.077 µM $ZnSO_4$, 0.011 µM $MoO_3$; pH 5.7 buffered with 5mM MES) once a week. Leaf tissue was harvested for ICP-MS analysis using a scalpel. Leaf samples were washed 3 times in MilliQ $H_2O$ to remove particles adhering to the outside before they were dried at 80°C for 24h. Root tissue was harvested by upending the contents of one pot into a large glass bowl. Roots were scooped from between the loose soil using tweezers and placed into water very briefly. Swirling them in water for a few seconds removed most of the adhering soil. Roots were removed and placed into fresh water containing a few drops of hair conditioner. The mixture was sonicated for 5 seconds and the now clean roots were rinsed in fresh water for a few seconds once more. Finally, roots were frozen in liquid nitrogen. From start to finish the root harvesting was finished within 2 minutes.

Inductively Coupled Plasma-Mass Spectronomy

Sample preparation as described previously[11]. Dried plant material was digested with 1ml concentrated trace metal grade nitric acid Primar Plus (Fisher Chemicals) spiked with Indium as an internal standard. Samples were heated in dry block heaters (SCP Science; QMX Laboratories) at 115°C for 4h. After cooling, digested samples were diluted to 10ml with 18.2 MΩcm Milli-Q Direct water (Merck Millipore) and subsequently analyzed using an ICP-MS, PerkinElmer NexION 2000 equipped with Elemental Scientific Inc. autosampler, in the collision mode (He). Twenty-four elements were monitored including the following stable isotopes: 7Li, 11B, 23Na, 24Mg, 31P, 34S, 39K, 43Ca, 48Ti, 52Cr, 55Mn, 56Fe, 59Co, 60Ni, 63Cu, 66Zn, 75As, 82Se, 85Rb, 88Sr, 98Mo, 111Cd, 208Pb and 115In. Liquid reference material composed of pooled samples was prepared before the beginning of a sample run and was used throughout the whole sample run. It was run after every ninth sample to correct for variation within ICP-MS analysis run. The calibration standards (with indium internal standard and blanks) were prepared from single element standards (Inorganic Ventures; Essex Scientific Laboratory Supplies Ltd, Essex, UK) solutions. Sample concentrations were calculated using external calibration methods within the instrument software. The final elements concentrations were obtained by normalizing the elements concentrations to the samples dry weight.

Phenotyping

Root hairs were analyzed from microscope pictures taken with a Zeiss StemiSV6 stereomicroscope with low magnification. Length and frequency was measured via ImagJ 1mm above the root tip and 2cm below the hypocotyl. The cell size was assessed using Fiji [13] to quantify microscope images. Cells were measured along the length of the root.

Confocal microscopy

Plants were grown on agar solidified ½ MS medium for 5 days, then stained in 6 µg/ml propidium iodide for 2 min and mounted on slides using cover slips No 1.5. A Leica SP5 Confocal microscope was used to image roots.

Confirmation of T-DNA insertions

Leaves were used for gDNA extraction. Material was frozen, pulverized and 500 µl extraction buffer (200 mM Tris/HCl pH 7.5, 250 mM NaCl, 25 mM EDTA, 0.5 % SDS) were added. DNA contained in the aqueous phase was precipitated using isopropanol (1:1) and washed with ethanol. After resuspension in 50 µl deionized water, gDNA was used for PCR amplification. Primers for amplification of DNA fragments around the T-DNA insertion sites were designed and plants tested for homozygous insertion of the respective T-DNA.

Analysis of transcript levels

Plant material was frozen in liquid nitrogen and RNA extracted by adding 1 ml of TRIzol® (Invitrogen) to 100 mg of pulverized material. After mixing 200 µl chloroform were added. After renewed shaking, cell debris was separated through centrifugation (12000g, 4°C, 15 min) and RNA in the liquid phase was precipitated by addition of equal volumes of Isopropanol. After centrifugation (12000g, 4°C, 10min) the liquid phase was discarded and the pelleted RNA washed with 70% ethanol before solubilizing it in 100µl RNAse-free water. Further purification was achieved by using the RNAeasy Kit from Quiagen, following the instruction manual of the manufacturer. Purity and concentration were analyzed via Nano-drop. For RNAseq analysis of RNA from plants grown on agar-solidified ¼ Hoagland grown plants the RNA quality was confirmed via RIN values (Agilent Tapestation System) and subsequently sent for sequencing to BGI Hong Kong. Single end (SE) sequencing was performed on a BGISEQ-500 RS Platform. Library preparation, sequencing, alignment and bioinformatics were performed by BGI according to their standard procedures. In short mRNA was enriched using Oligo(dT) magnetic beads, fragmented and reverse transcribed to double-strand cDNA by N6 random primers. The double stranded cDNA was then end repaired with phosphate at the 5' end and stickiness 'A' at the 3' end, and then ligated to adaptors with sticky 'T'. Using two specific primers the ligation product was amplified and the PCR product was cyclized by splint oligo and DNA ligases. The sequencing reactions were then performed on the library employing a 50SE strategy. The platform used was a benchtop high throughput sequencer BGISEQ-500 yielding 24,017,754 raw sequencing reads and then 24,007,388 clean reads after filtering low quality. After filtering, clean reads were mapped to reference genome (TAIR10 [14]) using HISAT and Bowtie2. The average mapping ratio with the reference genome was calculated to be 91.18%. Sequencing saturation and read randomness were checked for each sample before proceeding to ensure good quality. Gene expression as Fragments Per Kilobase of transcript per Million mapped reads (FPKM) was then calculated using the software package RSEM. This quantification tool computes the maximum likelihood abundance estimates by applying the statistical algorithm Expectation Maximization (EM). This includes paird-end (PE), variable-length reads, fragment length distributions and quality scores modelling to determine which transcripts are isoforms of one gene. The formula to calculate FPKM is: $FPKM = \frac{10^6 C}{NL/10^3}$. The benefit of this method is to eliminate the influence of differences in length as well as sequencing differences between samples on the gene expression calculation. Finally, differentially expressed genes were extrapolated by use of the NOISeq method. First, the noise distribution is modelled by calculating the log2 fold changes (M) from each samples gene expression and the absolute different value (D) of all pair conditions: $M^i = log_2(\frac{x_1^i}{x_1^i})$ and $D^i = |x_1^i - x_2^i|$. Then for each gene, the fold change is determined from the average expression in 2 different groups calculated from the three replicates and absolute different values for each gene will be calculated: $M_A = log_2(\frac{sample1\ avg}{sample2\ avg})$ and $D_A = |sample1\ avg - sample\ 2\ avg|$. $M_A$ and $D_A$ have to be different from the noise distribution model for the gene to be considered a differentially expressed gene (DEG). DEGs were then filtered for a fold change ≥2 and a diverge probability ≥0.8.

For RNAseq analysis of RNA from plants grown on agar solidifed ½ MS or soil RNA was isolated using the protocol from Paredes *et al.*, 2018 [15]. RNA was send for sequencing at Deep Seq University of Nottingham. In brief, RNA concentrations were measured using the Qubit Fluorometer and the Qubit RNA BR Assay Kit (ThermoFisher Scientific; Q10211) and RNA integrity was assessed using the Agilent TapeStation 4200 and the Agilent RNA ScreenTape Assay Kit (Agilent; 5067-5576 and 5067-5577). For each sample cDNA was generated from 200ng of total

RNA using the QuantSeq 3' mRNA-Seq library prep kit (Lexogen; 5001-5004). Indexed sequencing libraries were prepared using the Lexogen i7 6nt Index Set (Lexogen; 7001-7096). Libraries were quantified using the Qubit Fluorometer and the Qubit dsDNA HS Kit (ThermoFisher Scientific; Q32854). Library fragment-length distributions were analyzed using the Agilent TapeStation 4200 and the Agilent High Sensitivity D1000 ScreenTape Assay (Agilent; 5067-5584 and 5067-5585). Libraries were pooled in equimolar amounts and final library quantification performed using the KAPA Library Quantification Kit for Illumina (Roche; KK4824). The library pool was sequenced on the Illumina NextSeq 500 using a NextSeq 500 High Output Kit v2.5 75 cycle kit (Illumina; 20024906), to generate approximately 5 million 75bp single-end reads per sample. The raw data were imported into Galaxy Europe [16] and Salmon was used to quantify reads [17] and 3D RNAseq [18] used to filter, normalize and qc data, and to calculate DEGs. The package is able to calculate splice variants but only differential expression was used for this analysis since libraries were prepared such that only the 3' end was transcribed. Read counts and transcript per million reads (TPMs) were generated using tximport R package version 1.10.0 and scaledTPM method [19] with inputs of transcript quantifications from Salmon[17]. Low expressed genes were filtered based on analysing the data mean-variance trend. The expected decreasing trend between data mean and variance was observed when expressed transcripts were determined to have ≥ 6 of the 90 samples with count per million reads (CPM) ≥ 2, which provided an optimal filter of low expression. A gene was expressed if any of its transcripts with the above criteria was expressed. The TMM method was used to normalize the gene and transcript read counts to $log_2$-CPM (Bullard et al., 2010). A principal component analysis (PCA) plot showed the RNA-seq data did not have distinct batch effects. For differentially expressed genes, the $log_2$ fold change ($L_2$ FC) of gene/transcript abundance were calculated based on contrast groups and significance of expression changes were determined using t-test. P-values of multiple testing were adjusted with BH to correct false discovery rate (FDR) [20]. A gene was significantly differentially expressed in a contrast group if it had adjusted p-value < 0.05 and $L_2$ FC ≥ 0.5.

<u>Gene selection for Heatmap and PC analysis</u>

To find patterns of expression Thalemine was searched using the term "potassium transporters" which yielded 572 genes. PC analysis using the full set of genes lead to a significant correlation only in PC4 (Fdr adj. p value of r2 0.56). To enhance the sensitivity of the analysis a set of 31 genes with the highest Thalemine score were selected. 3 of these genes showed no expression. The remaining 28 were used for a PC and cluster analysis (Figure 1).

Generation of neo-tetraploid lines

The previously established protocols were modified and combined. Seeds of diploid progenitors were sown on Jiffies and germinated in long day conditions. 20 seedling per genotype, at the 2-leaf stage were treated with Colchicine (0.25% w/v in $H_2O$). One drop was placed on the shoot meristem. Contrastingly to the original protocol this was carried out on plants germinated on soil instead of plates [21]. By circumventing transferring plants to soil after their colchicine treatment the stress of the transfer can be avoided (Figure S14). Additionally, the workload is reduced. Plants which have been treated with colchicine stall in their development and only after 7 to 10 days it becomes apparent which of the shoot apexes will recover from the treatment. Surviving plants were grown to seed maturity. Seeds were sown and the germinating seedlings were pre-screened for increased trichome branching [21] and grown for 5 weeks (Figure S14) under short day conditions. At this stage one leaf was harvested and tested for whole genome duplication (WGD) using flow cytometry. The isolation protocol was adapted [22,23] for high throughput. In this manner the isolation and analysis of 60 samples can be achieved in one day. The leaves were mechanically disrupted by peeling the lower leaf epidermis using tape [23]. The exposed leaves were shaken in 600µl extraction buffer (15mM HEPES, 1mM Na-EDTA, 80mM KCl, 20mM NaCl, 200mM Sucrose, 0.2% Trition-X, 0.5mM Spermin, pH 6.1), 0.3µl RNase (100ng/µl) was added and samples were incubated at RT for 10min. 10µl of PI (1mg/ml) was added and samples stored at 4°C, in the dark for at least 1h until analysis. 400µl of each sample were transferred to analysis tubes and run on the DB FACS Canto. Settings for the machine: FSC 376V, SSC 330V, PI 369V, cut-off 35000, events 10 – 50 thousand. On a $logSSC_{area}$ vs $logPI_{area}$ plot distinct fractions were gated and displayed in a $PI_{area}$ histogram. Data were analyzed using Kaluza software (Version 2.1.00001.20653). Ploidy was determined by the highest peak from the histogram, diploid and tetraploid Col-0 nuclei extracts were used to identify 2x and 4x peaks. Plants analyzed via FACS were transferred to long day conditions and further grown until seed maturity. Seeds harvested from these plants were collected for phenotypic analysis of neo-tetraploid mutant lines. Nomenclature: Diploid mutant lines were described as per general conventions ie. *hak5* gene, line *hak5-3* referring to line N574868 from the NASC stock collection or SALK_074868.54.75.x. Individual colchicine treated plants were given consecutive number ie.

*hak5-3_1* followed by *hak5-3_2* etc. Lines derived from these independent colchicine treated plants were annotated with #consecutive numbers ie. *hak5-3_1#1* or *hak5-3_2#1* represent two independent duplication events of the same diploid progenitor. Additionally, chromosomal spreads of floral buds of plants selected via flow cytometry were prepared [24] and chromosomes were counted (Figure S14). Neo-tetraploid nuclei contain 20 chromosomes which were visible during the metaphase or first condensation during prophase (Figure S14).

Statistical analysis

Figures were created and statistical analyses were performed with R version 4.1.0 (2021-05-18)[25] using RStudio Version 1.0.143 (www.rstudio.com) [26]. Platform: x86_64-w64-mingw32/x64 (64-bit), Running under: Windows 10 x64 (build 19043). The following packages were used: PCAtools (version 2.4.0), ggrepel (version 0.9.1), nsprcomp (version 0.5.1-2), factoextra (version 1.0.7), FactoMineR (version 2.4), RFLPtools (version 1.9), clValid (version 0.7), pheatmap (version 1.0.12), heatmaply (version 1.2.1), plotly (version 4.9.4.1), dendextend (version 1.15.1), veccompare (version 0.1.0), VennDiagram (version 1.6.20), futile.logger (version 1.4.3), scales (version 1.1.1), kableExtra (version 1.3.4), knitr (version 1.33), flextable (version 0.6.7), corrplot (version 0.90), Hmisc (version 4.5-0), Formula (version 1.2-4), devtools (version 2.4.2), usethis (version 2.0.1), ggpmisc (version 0.4.0), ggpp (version 0.4.1), gplots (version 3.1.1), cowplot (version 1.1.1), sjPlot (version 2.8.9), bestNormalize (version 1.8.0), multcompView (version 0.1-8), multcomp (version 1.4-17), TH.data (version 1.0-10), MASS (version 7.3-54), survival (version 3.2-11), mvtnorm (version 1.1-2), emmeans (version 1.6.2-1 ggpubr (version 0.4.0), cluster (version 2.1.2), viridis (version 0.6.1), viridisLite (version 0.4.0), RColorBrewer (version 1.1-2), gridExtra (version 2.3), tidyquant (version 1.0.3), quantmod (version 0.4.18), TTR (version 0.24.2), PerformanceAnalytics (version 2.0.4), xts (version 0.12.1), zoo (version 1.8-9), lubridate (version 1.7.10), ggfortify (version 0.4.12), rcompanion (version 2.4.1), forcats (version 0.5.1), stringr (version 1.4.0), dplyr (version 1.0.7), purrr (version 0.3.4), readr (version 2.0.0), tidyr (version 1.1.3), tibble (version 3.1.3), ggplot2 (version 3.3.5), tidyverse (version 1.3.1), Rmisc (version 1.5), plyr (version 1.8.6), lattice (version 0.20-44).

Gene ontology (GO) enrichments were calculated using the PANTHER tool (http://www.pantherdb.org/). A statistical overrepresentation test using the Fisher's Exact test and Bonferroni correction for multiple testing was used to calculated p-values. These take the group size for each GO term into account as well as the number of DEGs in total and the number of GO annotatable genes. The resulting value is thus scaled. Gene ontology terms with $p \leq 0.05$ were selected for interpretation. The hierarchy display was used to select the most detailed group amongst all related GO terms and results displayed for this group only. The full results can be found in Table S2. A GO enrichment for other contrast groups besides root expression in diploid vs neo-tetraploid wild type, which is shown in the main text, can be found in Figure S12-S13.

**Supplementary Figures**

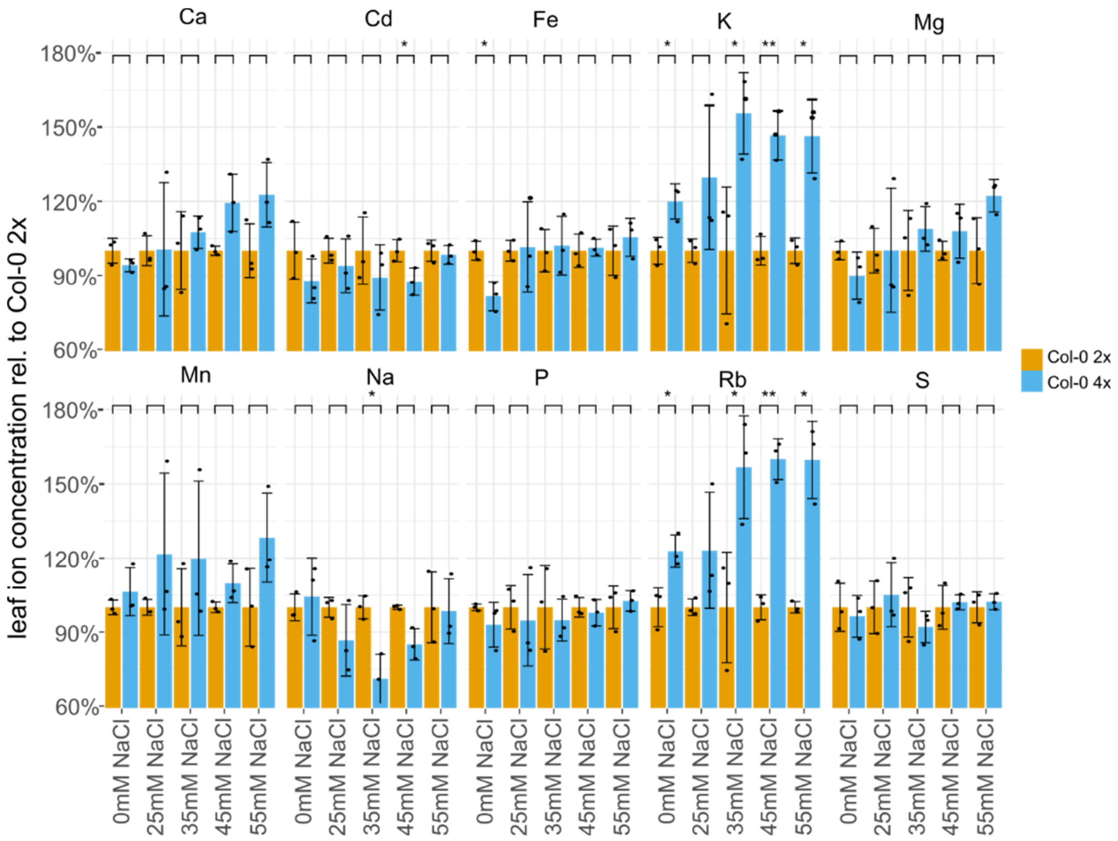


**Figure S1: Ploidy dependent ionomic differences**. The leaf ionome of diploid and neo-tetraploid *A. thaliana* wild type (Col-0) plants was assessed in response to varying salt stress. The results show genotype and treatment dependent changes. Plants were grown on agar solidified ¼ Hoagland medium for 7 days. n= 3-6, Different letters show significant differences between genotypes and treatments but within elements, the distinctions are based on a 2-way ANOVA for ploidy and treatment with Posthoc Tukey test. Yellow dot: averages. Diploid (2x), neo-tetraploid (4x)

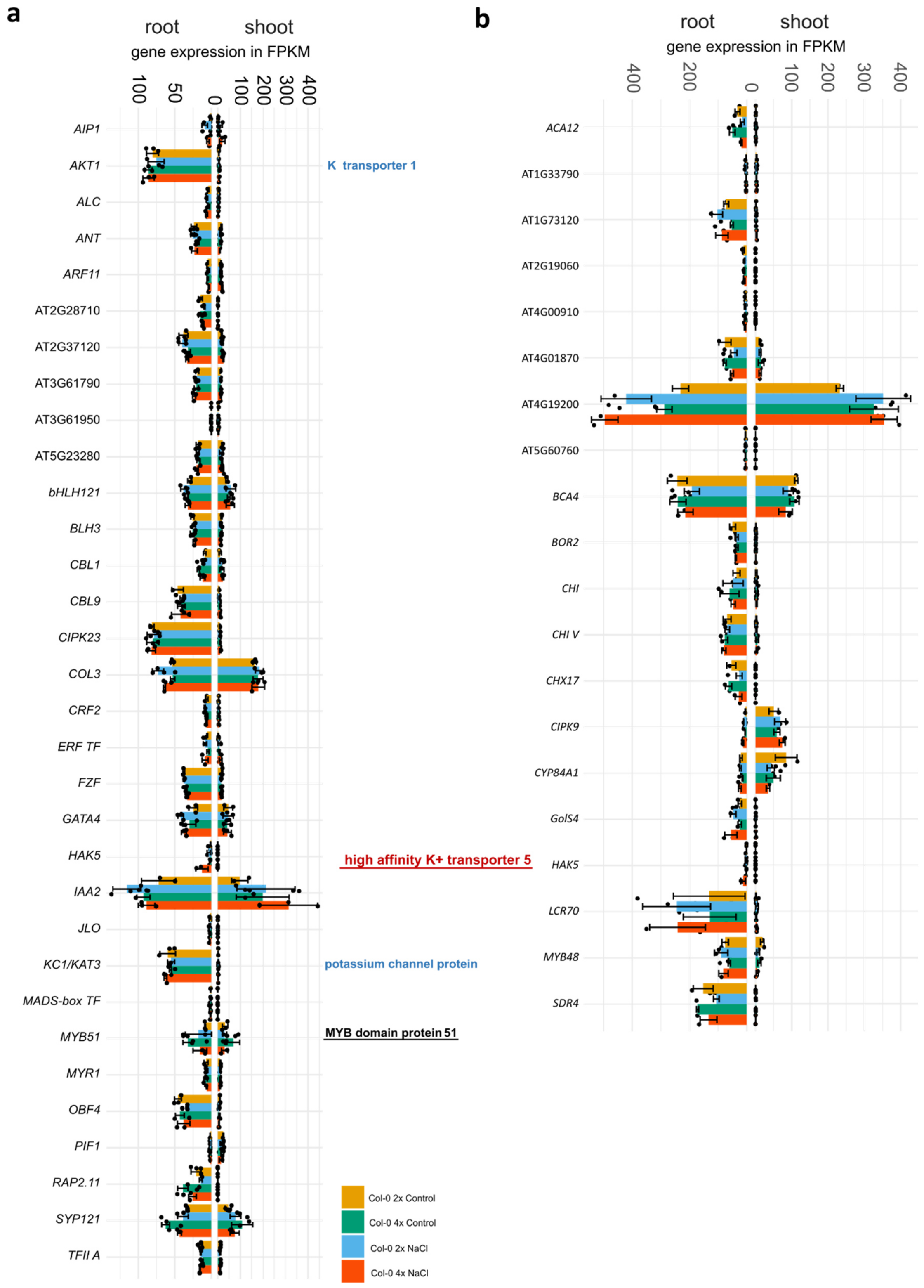


**Figure S2: Expression of Low K signalling in neo-tetraploids**. The expression of components of the low K signalling machinery as defined by **a)** Nieves-Cordones *et al*., 2014 and Hong et al., 2013

[27,28] and by **b)** a cross comparison of K starvation genes identified in a gene expression study by Forieri *et al*., 2016, Gierth *et al*., 2005 and Hampton *et al*., 2004[29–31]. Bold gene descriptions mark genes for which an induction by low K has been seen as well as a physical interaction with the promotor of *HAK5*. In red and blue are genes for the high affinity and low affinity K uptake system respectively and in. Underlined are genes defined as DE between diploids and neo-tetraploids. With * are genes defined as DE after Na-stress. n=3+/-se. Diploid (2x), neo-tetraploid (4x)

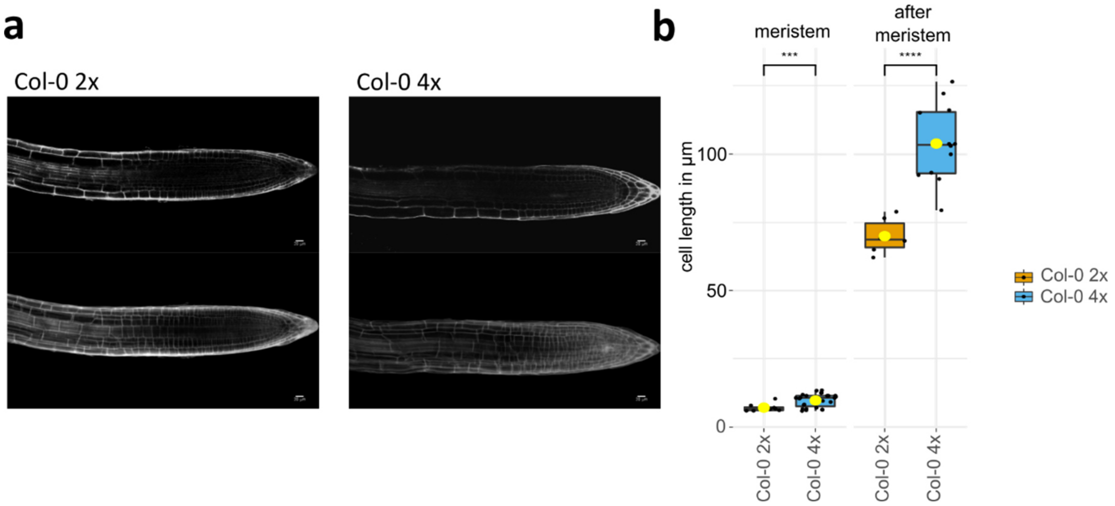


**Figure S3: Cell size of neo-tetraploids. a)** Confocal image of propidiumiodid stained roots of diploid and neo-tetraploid plants grown on ½ MS, agar solidified, media. The size of meristematic and fully expanded cells were measured. **b)** General cell size difference in wild type Col-0 neo-tetraploids. Cells of different types are larger. The box plot shows the quantification of microscope images. n=6-26, Pairwise comparison using t-test indicates significant differences between diploids and neo-tetraploids. p-value ≤: *0.05, **0.01, ***0.001, ****0.0001. Diploid (2x), neo-tetraploid (4x)

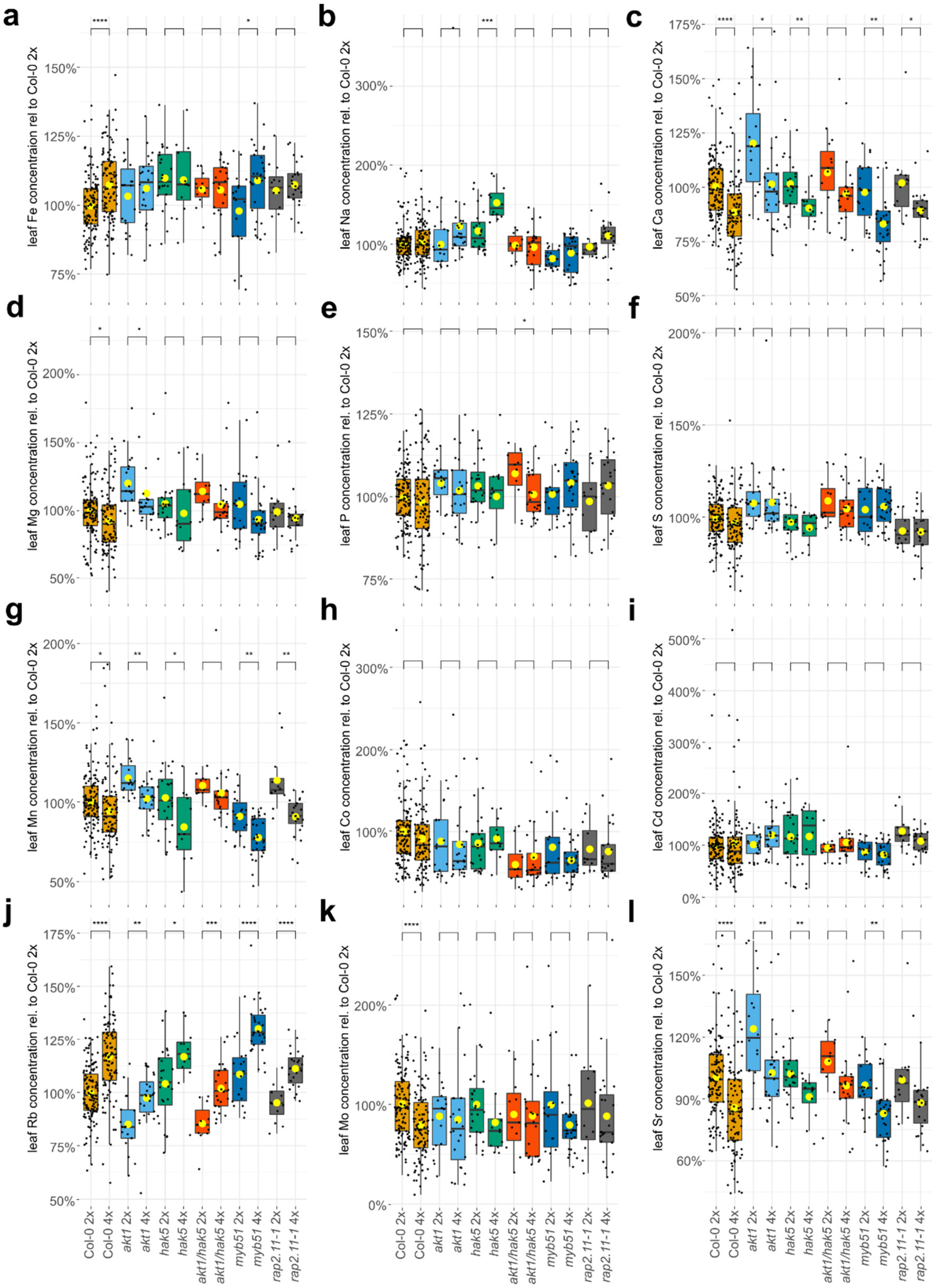
a
b
c
d
e
f
g
h
i
j
k
l
leaf Fe concentraion rel to Col-0 2x
leaf Na concentration rel. to Col-0 2x
leaf Ca concentration rel. to Col-0 2x
leaf Mg concentration rel. to Col-0 2x
leaf P concentration rel. to Col-0 2x
leaf S concentration rel. to Col-0 2x
leaf Mn concentration rel. to Col-0 2x
leaf Co concentration rel. to Col-0 2x
leaf Cd concentration rel. to Col-0 2x
leaf Rb concentration rel. to Col-0 2x
leaf Mo concentration rel. to Col-0 2x
leaf Sr concentration rel. to Col-0 2x
Col-0 2x
Col-0 4x
akt1 2x
akt1 4x
hak5 2x
hak5 4x
akt1/hak5 2x
akt1/hak5 4x
myb51 2x
myb51 4x
rap2.11-1 2x
rap2.11-1 4x

**m**

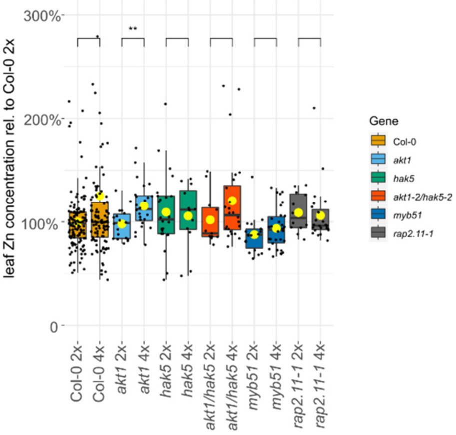


**Figure S4: Ploidy dependent ionomic differences**. **a-n)** The leaf ionome of diploid and neo-tetraploid *A. thaliana* wild type (Col-0) plants was assessed. The results show genotype dependent changes. Plants were grown on peat-based soil for 5 weeks. n= 11-121, Pairwise comparison using t-test indicates significant differences between diploids and neo-tetraploids. p-value ≤: *0.05, **0.01, ***0.001, ****0.0001 Yellow dot: averages. All elements within the limit of quantification (LOQ) were plotted. Diploid (2x), neo-tetraploid (4x)

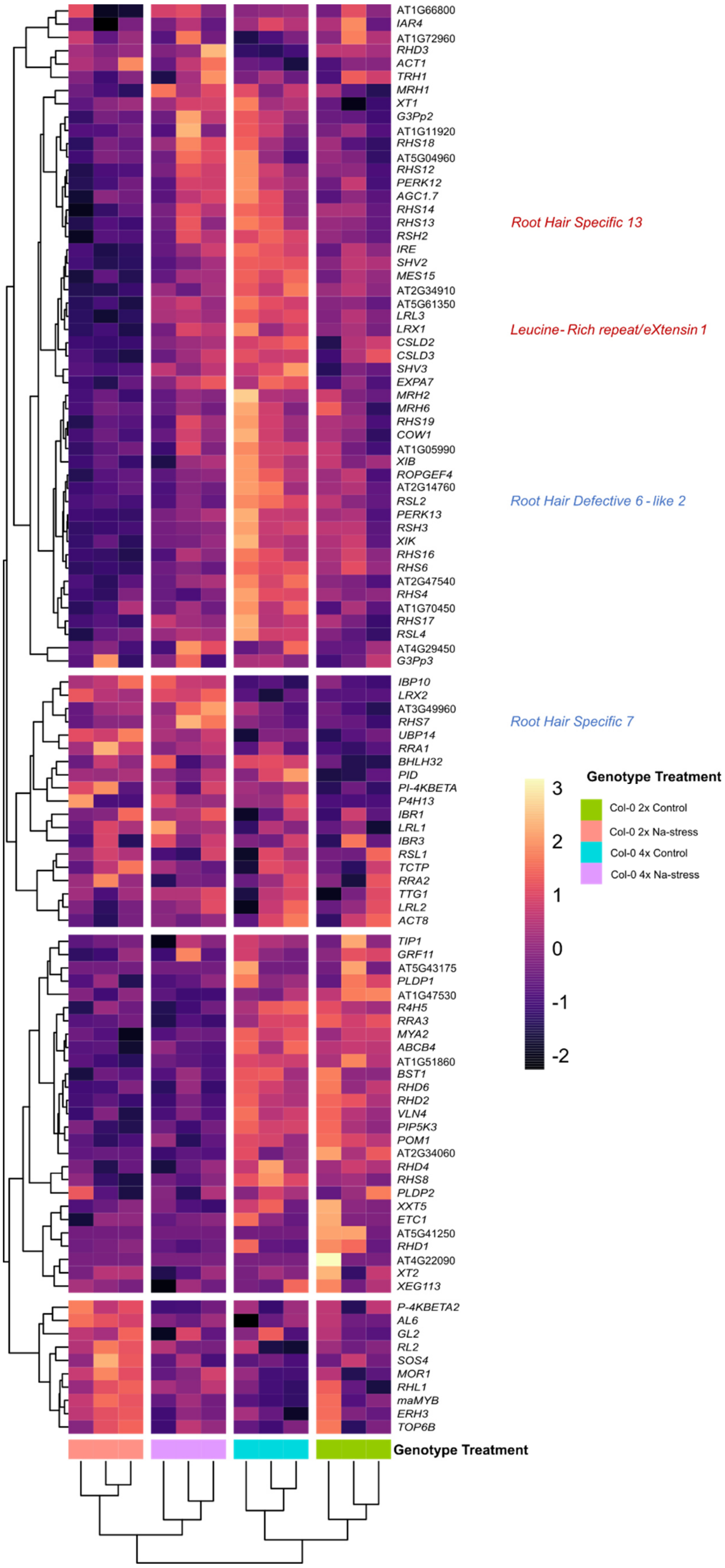
AT1G66800
IAR4
AT1G72960
RHD3
ACT1
TRH1
MRH1
XT1
G3Pp2
AT1G11920
RHS18
AT5G04960
RHS12
PERK12
AGC1.7
RHS14
RHS13
RSH2
IRE
SHV2
MES15
AT2G34910
AT5G61350
LRL3
LRX1
CSLD2
CSLD3
SHV3
EXPA7
MRH2
MRH6
RHS19
COW1
AT1G05990
XIB
ROPGEF4
AT2G14760
RSL2
PERK13
RSH3
XIK
RHS16
RHS6
AT2G47540
RHS4
AT1G70450
RHS17
RSL4
AT4G29450
G3Pp3
IBP10
LRX2
AT3G49960
RHS7
UBP14
RRA1
BHLH32
PID
PI-4KBETA
P4H13
IBR1
LRL1
IBR3
RSL1
TCTP
RRA2
TTG1
LRL2
ACT8
TIP1
GRF11
AT5G43175
PLDP1
AT1G47530
R4H5
RRA3
MYA2
ABCB4
AT1G51860
BST1
RHD6
RHD2
VLN4
PIP5K3
POM1
AT2G34060
RHD4
RHS8
PLDP2
XXT5
ETC1
AT5G41250
RHD1
AT4G22090
XT2
XEG113
P-4KBETA2
AL6
GL2
RL2
SOS4
MOR1
RHL1
maMYB
ERH3
TOP6B
Root Hair Specific 13
Leucine- Rich repeat/eXtensin 1
Root Hair Defective 6 - like 2
Root Hair Specific 7
3
2
1
0
-1
-2
Genotype Treatment
Col-0 2x Control
Col-0 2x Na-stress
Col-0 4x Control
Col-0 4x Na-stress
Genotype Treatment

**Figure S5: Neo-tetraploids and root hair genes.** Heat map shows expression of root hair specific genes in diploid and neo-tetraploid wild type (Col-0) roots under control conditions and Na-stress. n=3 +/- SE. In blue are DEG responsive to Na-stress only in neo tetraploids. In red are genes with significant expression differences in neo-tetraploids Vs diploids at Na-stress. Diploid (2x), neo-tetraploid (4x)

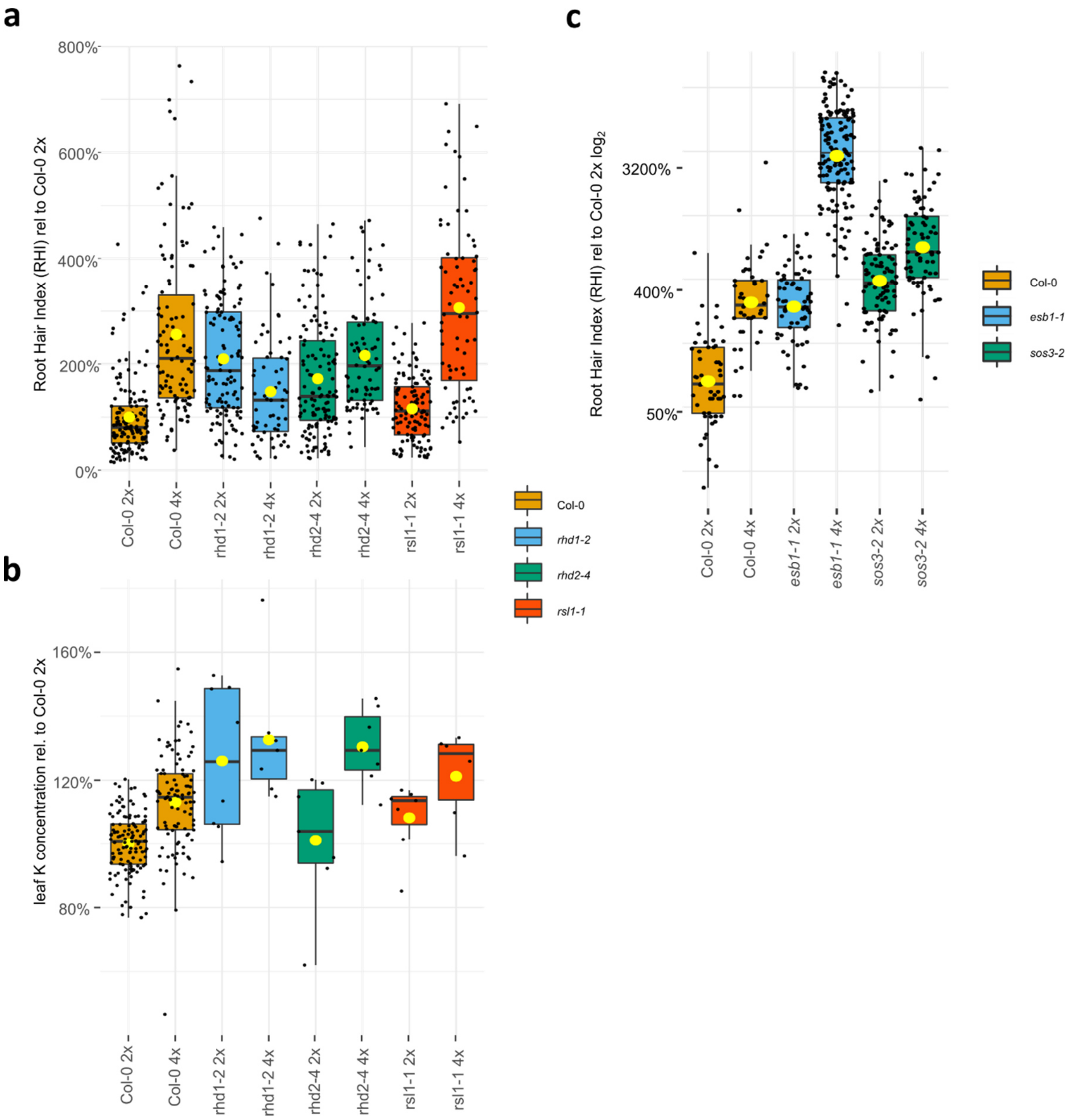


**Figure S6: Neo-tetraploids and their root hairs. a)** Boxplot shows differences in root hair index (RHI) for diploid and neo-tetraploid lines. n=51-134, A two-way ANOVA with post hoc Tukey test showed significant effects for both ploidy (p≤0) and genotypes *rhd1-2* and *rhd2-4* (p≤0 for both) and *rsl1-1* (p≤0.01) as well as for the ploidy*genotype (**rhd1-2* and *rhd2-4*) interaction (p≤0 for both). None of the mutants show a strong reduction in RHI as could be seen for the *rhd6-3/rsl1-1* and *rsl4-1* both of which suppress the ploidy K phenotype. **b)** Boxplot shows leaf K concentration of plants assessed for their RHI in A. n= 6-109. A two-way ANOVA with post hoc Tukey test showed significant effects for both ploidy (p≤0) and genotypes *rhd1-2* (p≤0 ) but not for *rhd2-4* and *rsl1-1*. As well as for one of the ploidy*genotype (**rhd2-4*) interactions (p≤0.05). The neo-tetraploid root hair mutants do not suppress the K phenotype. **c)** Boxplot shows log2 RHI for wild type, *esb1-1* and *sos3-2* diploids and neo-tetraploids. n=42-142, A two-way ANOVA with post hoc Tukey test showed significant effects for both ploidy (p≤0) and genotypes *esb1-1* and *sos3-2* (p≤0 for both) as well as for the ploidy*genotype interaction (p≤0 for both). Both mutants, which suppress the K phenotype do show root hairs at least as long/dense as the wild type and also display the typical

RHI increase after WGD. It can therefore be excluded that a loss of root hairs is the explanation for the loss of the K phenotype in neo-tetraploid *esb1-1* and *sos3-2* as it is for *rhd6-3/rsl1-1*. Yellow dot: averages. Diploid (2x), neo-tetraploid (4x)

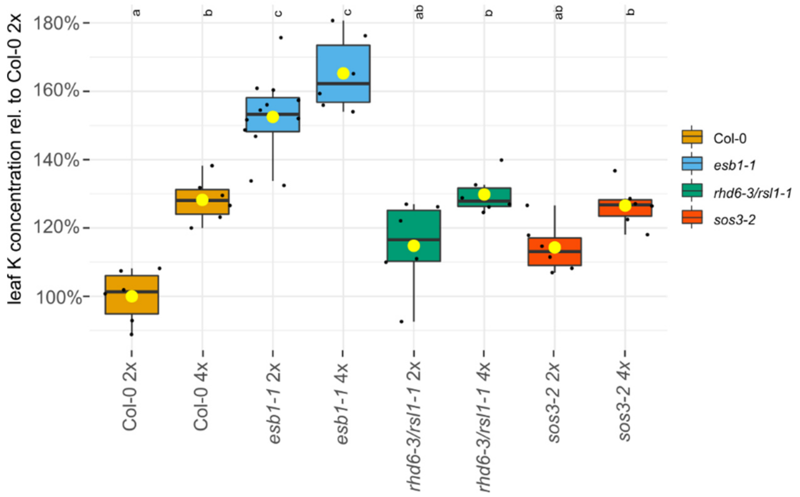


**Figure S7: Leaf K for soil grown plants used for RNAseq.** Boxplots show the K concentration relative to the leaf K content of diploid wild type. n= 6. Different letters show significant differences between groups, the distinctions are based on a 2-way ANOVA with Posthoc Tukey test. Diploid (2x), neo-tetraploid (4x)

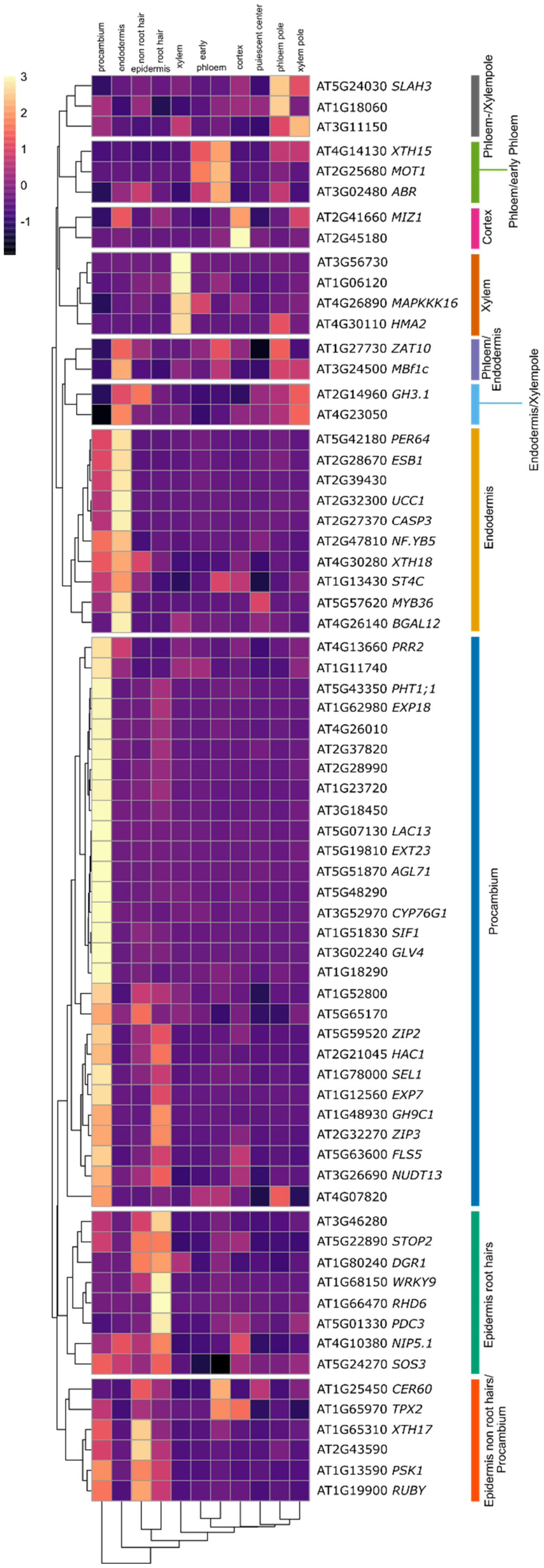
3
2
1
0
-1
procambium
endodermis
non root hair
root hair
epidermis
xylem
early
phloem
cortex
quiescent center
phloem pole
xylem pole
AT5G24030 SLAH3
AT1G18060
AT3G11150
AT4G14130 XTH15
AT2G25680 MOT1
AT3G02480 ABR
AT2G41660 MIZ1
AT2G45180
AT3G56730
AT1G06120
AT4G26890 MAPKKK16
AT4G30110 HMA2
AT1G27730 ZAT10
AT3G24500 MBf1c
AT2G14960 GH3.1
AT4G23050
AT5G42180 PER64
AT2G28670 ESB1
AT2G39430
AT2G32300 UCC1
AT2G27370 CASP3
AT2G47810 NF.YB5
AT4G30280 XTH18
AT1G13430 ST4C
AT5G57620 MYB36
AT4G26140 BGAL12
AT4G13660 PRR2
AT1G11740
AT5G43350 PHT1;1
AT1G62980 EXP18
AT4G26010
AT2G37820
AT2G28990
AT1G23720
AT3G18450
AT5G07130 LAC13
AT5G19810 EXT23
AT5G51870 AGL71
AT5G48290
AT3G52970 CYP76G1
AT1G51830 SIF1
AT3G02240 GLV4
AT1G18290
AT1G52800
AT5G65170
AT5G59520 ZIP2
AT2G21045 HAC1
AT1G78000 SEL1
AT1G12560 EXP7
AT1G48930 GH9C1
AT2G32270 ZIP3
AT5G63600 FLS5
AT3G26690 NUDT13
AT4G07820
AT3G46280
AT5G22890 STOP2
AT1G80240 DGR1
AT1G68150 WRKY9
AT1G66470 RHD6
AT5G01330 PDC3
AT4G10380 NIP5.1
AT5G24270 SOS3
AT1G25450 CER60
AT1G65970 TPX2
AT1G65310 XTH17
AT2G43590
AT1G13590 PSK1
AT1G19900 RUBY
Phloem-/Xylempole
Phloem/early Phloem
Cortex
Xylem
Phloem/ Endodermis
Endodermis/Xylempole
Endodermis
Procambium
Epidermis root hairs
Epidermis non root hairs/ Procambium

**Figure S8: Heatmap used to display cell-type expression of 80 genes**. ePlant http://bar.utoronto.ca/eplant/ was used to obtain data on root cell type specific expression patterns for 80 selected genes. Data for 68 genes was available on the platform. Expression was normalized to "ROOT_CTRL" and averaged within procambium, endodermis, epidermis non root hair, epidermis root hair, xylem, cortex, early phloem, phloem, quiescent centre, phloem pole and xylem pole. A heatmap of the normalized average expression was generated and clustered. High expression is indicated by bright colours, low expression by dark colours. Clusters were used to annotate network analysis in Figure 4 using the colour scheme indicated next to the heatmap.

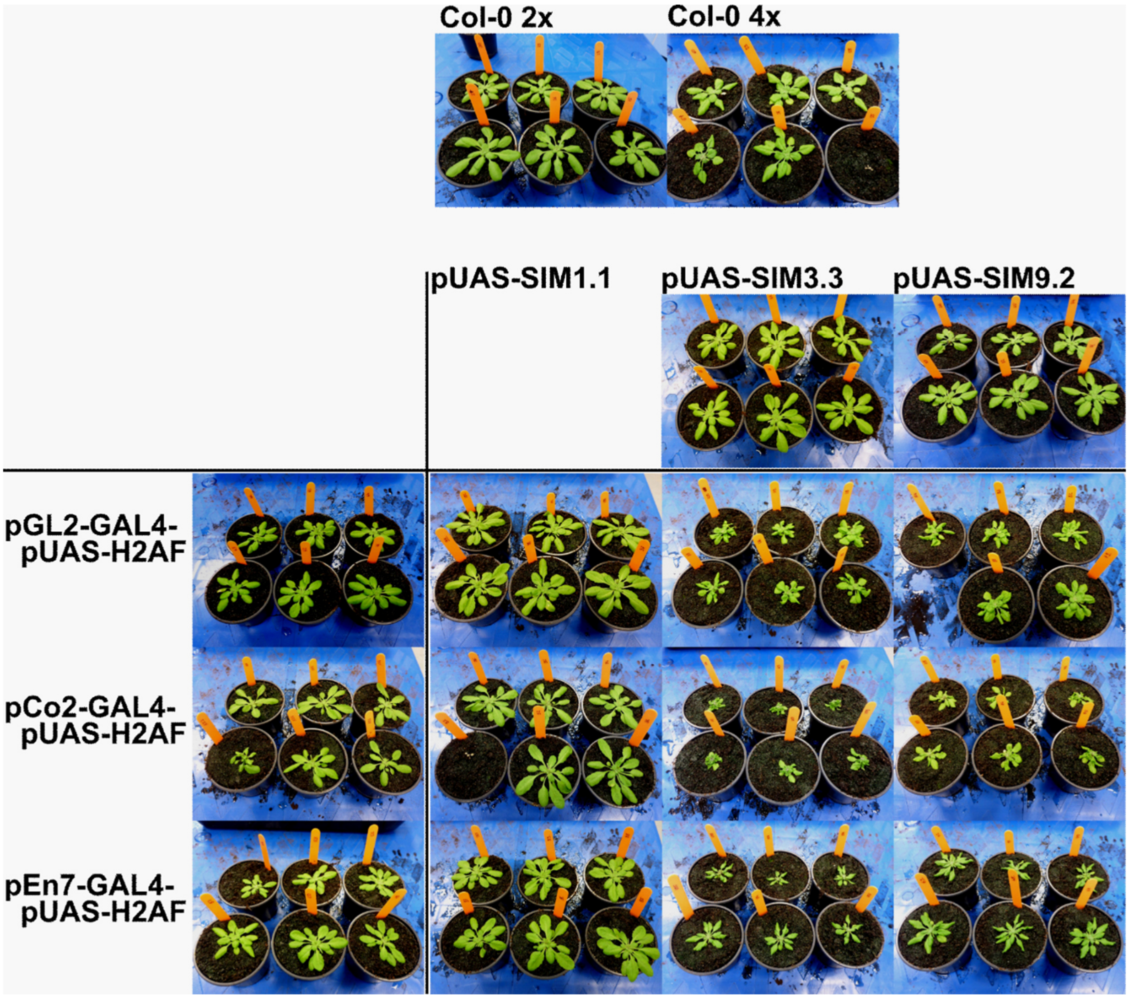


**Figure S9: SIM overexpression in Epidermis, Cortex and Endodermis.** Pictures of plants expressing *SIM* in a tissue specific manner shows an impact on plant growth and morphology [2]. Lines *1.1 X GL2/Co2/En7* are not affected due to silencing of the *SIM* construct.

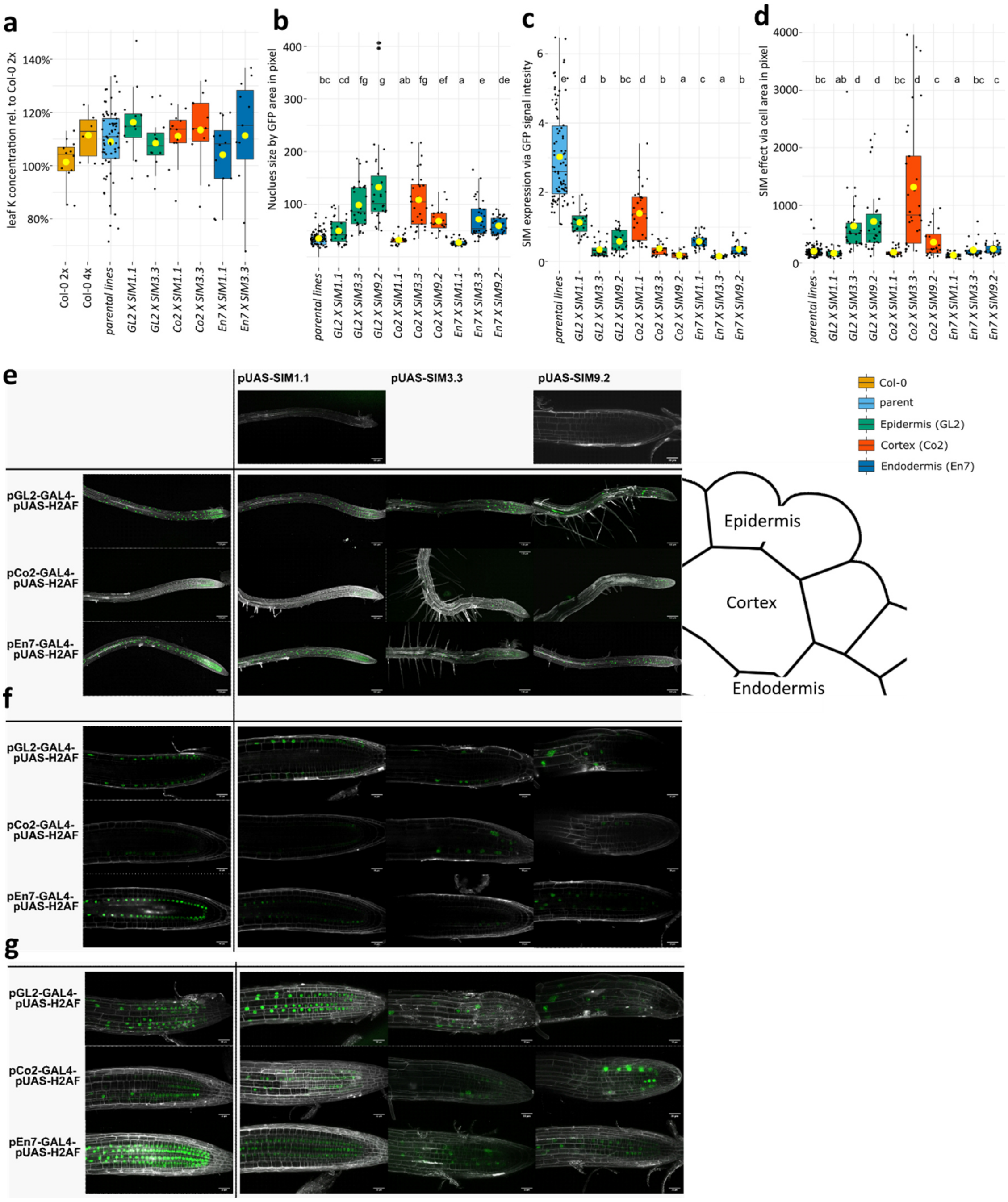


**Figure S10: Root Hairs after SIM overexpression.** Microscope images of roots of plants expressing GFP and (in F1 plants of crosses with *pUAS-SIM* lines) *SIM* in epidermis, cortex and endodermis. 5-day old plants, grown on ½ MS agar solidified plates containing no sucrose, are stained with propidiumiodid for 3min. **a)** Boxplots show the K concentration relative to the leaf K content of diploid wild type. n=6-65. Diploid (2x), neo-tetraploid (4x) Pairwise comparison using t-test indicates significant differences between parent and *SIM* expressing line for endodermal expression. p-value ≤: *0.05, **0.01, ***0.001, ****0.0001 Yellow dot: averages **b)** Quantification of the GFP signal allows an estimation of *SIM* expression which is higher in the line *En7 X SIM9.2* than *En7 X SIM3.3* and also in *GL2 X SIM9.2* than *GL3 X SIM3.3.* The signal intensity was adjusted to compensate for different gain. **c)** The effect of SIM on endoreduplication was assessed by measuring the GFP area which is correlated to the nuclei area. Nuclei of *GL2/Co2/En7 X 1.1* lines are smaller than those of *GL2/Co2/En7 X 9.2* and *3.3* lines, indicating that *SIM* was silenced in this cross, which serves as an additional control. **d)** The cell area in GFP expressing cells was measured. To avoid the impact of cell elongation cells of the root tips were chosen for this measurement. The effect of *SIM* is related to cell size as expected. n=10-30, letters show significant ($p<0.05$) differences between groups, result of a one-way ANOVA. **e)** Lower magnification shows the early development of root hairs in *GL2/Co2/En7 X SIM* lines. Images are maximum projections GFP gain varied in order to ensure visibility but avoid overexposure. Schematic indicates in which cell type *SIM* is expressed. **f)** Central region of the root at a higher magnification shows the localization of the GFP signal in the respective tissue. **g)** Maximum projection through the whole root shows size of nuclei in *GL2/Co2/En7 X 9.2* and *3.3* lines.

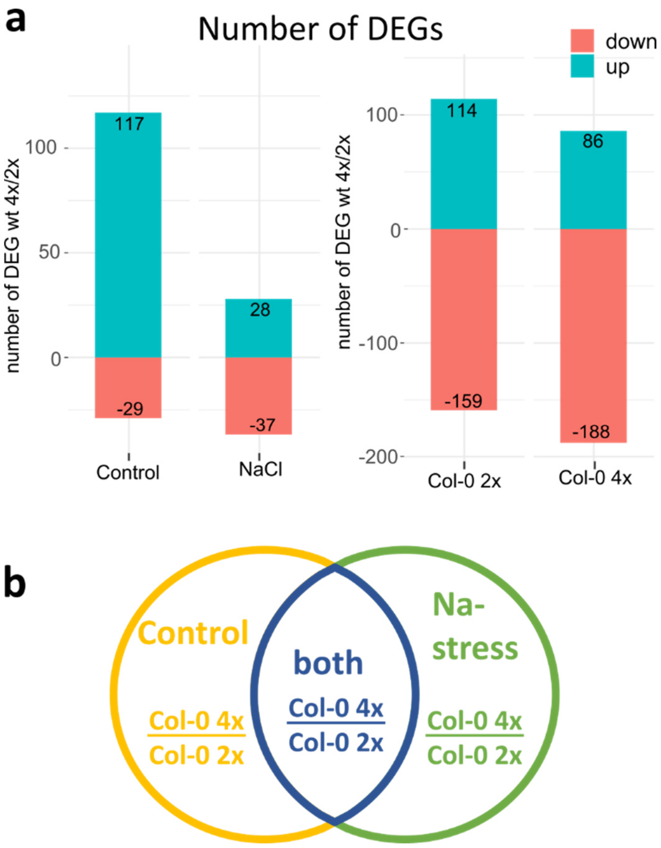


**Figure S11: RNAseq of diploid and neo-tetraploid wild type plants grown on ¼ Hoaglands, agar solidified medium . a)** Number of DEGs between either diploids and neo-tetraploids (under control and Na stress) or between control or Na stressed plants (either for diploids (2x) or for neo-tetraploids (4x)) are shown for root expressed genes. **b)** A Venn diagram shows the comparison between DE genes which was used for the GO analysis shown in main Figure 5A and in Figure S14. Genes were defined as being DE in neo-tetraploids under either control conditions (yellow) or Na stress (green) or both (blue). The colour scheme was used in Figure 5A and S14 to indicate which group of genes was used for the respective GO enrichment analysis. Diploid (2x), neo-tetraploid (4x)

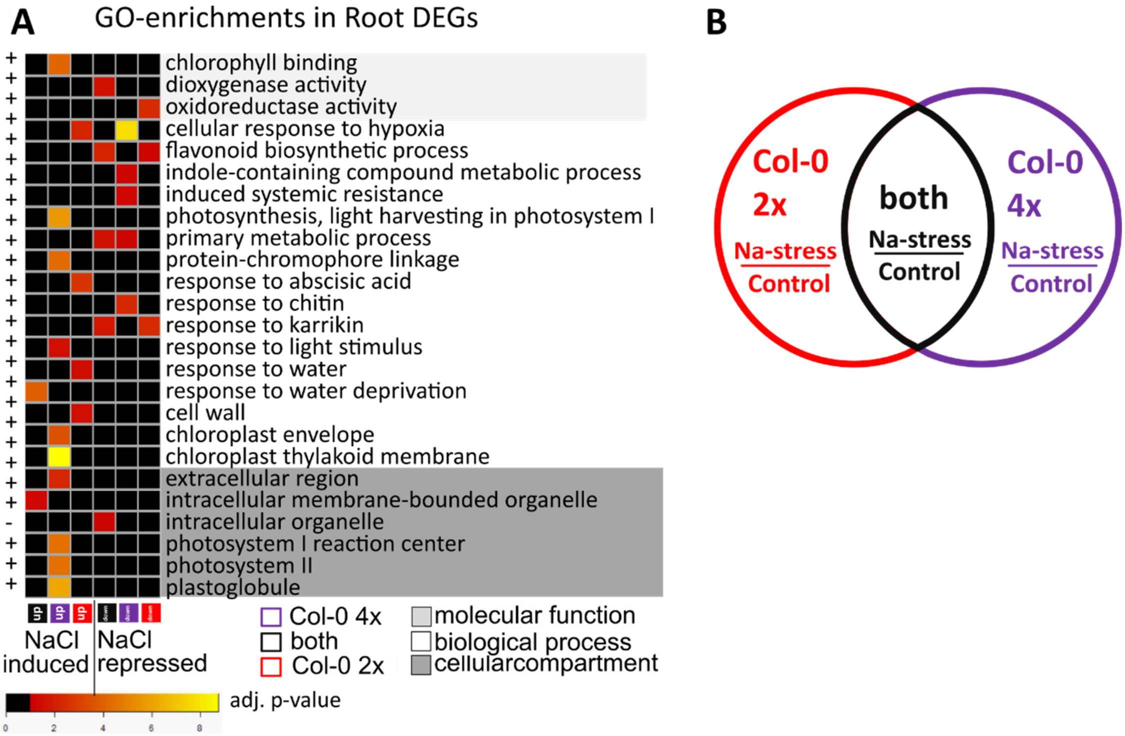


**Figure S12: PANTHER GO-enrichment analysis of the effect of Na on the root transcriptome. a)** The heatmap shows p-values of 0.05 or lower, with yellow representing more significant -log(10) adjusted p-values for GO groups enriched in at least one of the 6 groups of genes. Analysis was done using PANTHER. Displayed are only the most defined terms or child terms. A full list can be found in Table S2. Test Type: Fisher's Exact, Bonferroni correction for multiple testing, GO database release 2019-12-09. The groups of genes are annotated with different colours underneath the heatmap and the gene selection process is shown in **b)** A Venn diagram shows the comparison between DEGs which was used for the GO analysis shown in Figure S13 A. Genes were defined as being DEGs after Na stress in either diploids (red) or neo-tetraploids (purple) or both (black).

Abbreviations: *=oxidoreductase activity, acting on paired donors, with incorporation or reduction of molecular oxygen, **=chloroplast thylakoid membrane protein complex, transc.=transcription, resp.=response, membr.=membrane. Diploid (2x), neo-tetraploid (4x)

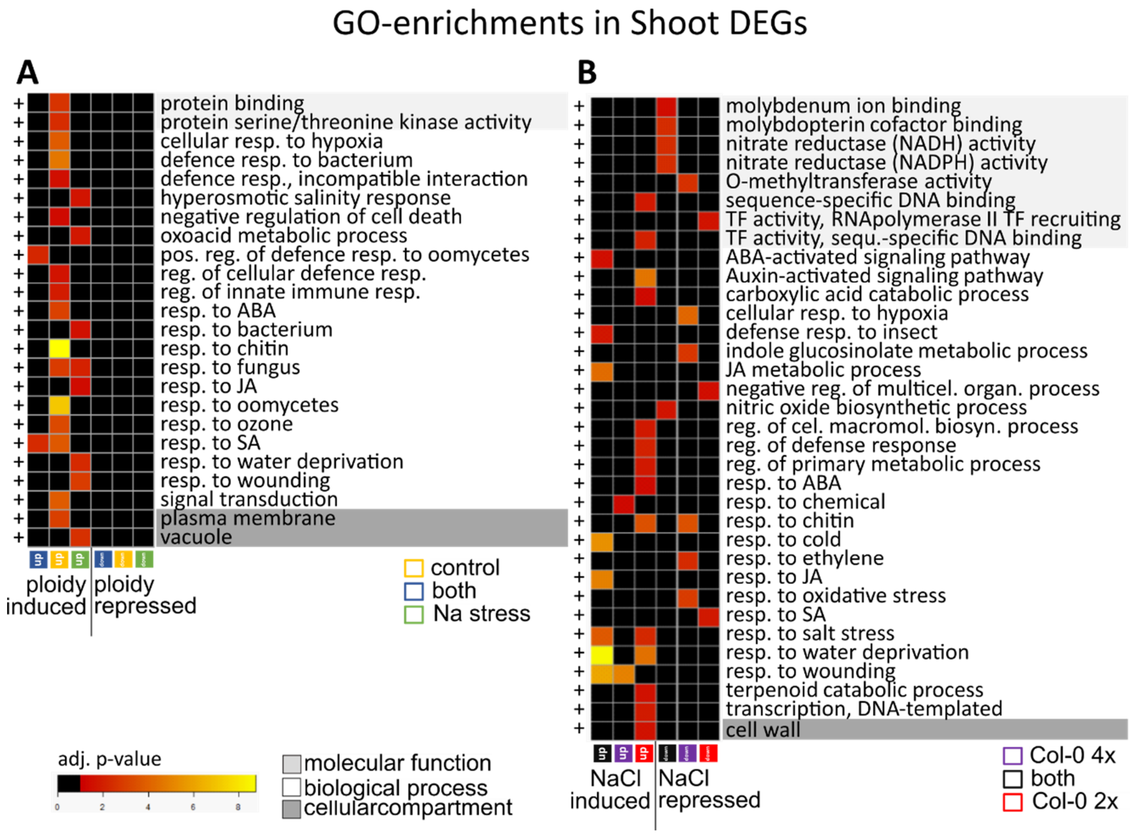


**Figure S13: PANTHER GO-enrichment analysis of shoot transcriptome. a-b)** The heatmaps shows p-values of 0.05 or lower, with yellow representing more significant -log(10) adjusted p-values for GO groups enriched in at least one of the 6 groups of genes. Analysis was done using PANTHER. Displayed are only the most defined terms or child terms. A full list can be found in Table S2. Test Type: Fisher's Exact, Bonferroni correction for multiple testing, GO database release 2019-12-09. The groups of genes are annotated with different colours underneath the heatmap and the gene selection process is shown in Figure S12B.

Abbreviations: transc.=transcription, resp.=response, membr.=membrane. Diploid (2x), neo-tetraploid (4x)

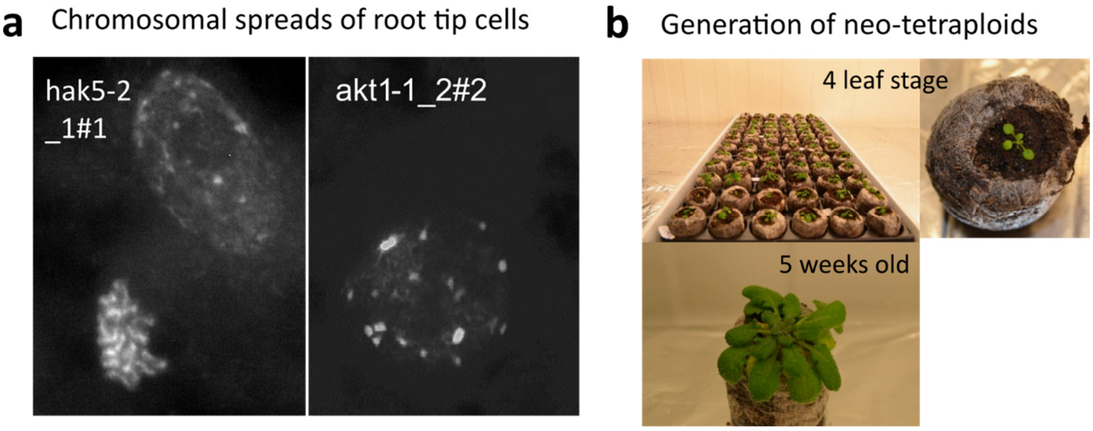


**Figure S14: Generation of new tetraploid lines. c)** To generate new 4x lines plants of 4-leaf-stage were treated with colchicine and seeds collected of surviving plants. Among the progeny those with 4-branched trichomes were selected, grow for 5 weeks at which point a leaf was harvested for nuclei isolation. Pictures show yiffi® growing plants in the 2-leaf stage just after colchicine treatment and a full setup of yiffies® containing a set of 8 lines each growing on 13 yiffies® in the WGD pipeline. Finally, a fully grown 5 week old plant in the 1st generation after WGD used to harvest nuclei before re-potting for seed generation. **a)** Nuclei, stained with PI were analyzed via flow cytometry. Profiles show the number of counts for PI signal of different intensities. Different colors mark different nuclei sizes, recognizable by more intense staining of nuclei with higher amounts of DNA. The diploid wild type and previously generated neo-tetraploid wild type [5] were used to calibrate the instrument settings and to detect the correct peaks. **b)** Microscopic images of chromosomal spreads show the 20 chromosomes of neo-tetraploid lines. This technique was used to verify the more high-throughput FACS analysis.

**Table S1.** Table S1: List of *A. thaliana* lines used and generated for this study. Whenever possible several alleles were used for each gene of interest. If not possible it was attempted to generate 2 or more independent neo-tetraploid lines. Nomenclature was such: New lines were given a number/letter code which can be found in the "Comment" column for the original T-DNA line. For example, 6 for N653619 or *sur1-8*. Each plant which survived colchicine treatment was given incremental number such as 6-1, 6-8 being the 1st and the 8th plant to survive colchicine. Two seeds from the survivors were germinated and given a #1 or #2 addition making it 6-1#1 and 6-1#2 and 6-8#1 and 6-8#2. These lines were assessed for ploidy and neo-tetraploid lines kept for experimentation.

**Table S2**: Full results of GO enrichment analysis of a subset of genes DE between wild type diploid and neo-tetraploids in roots either under control conditions, or under Na stress or both. See Figure S12B to see a visual representation of the gene selections. The most detailed group (child term) is displayed graphically in a heatmap Figure 5 and Figure S13-15 as is highlighted here in the column "most detailed term". Each group of GO groups hierarchically connected to each other is marked in alternating yellow/grey and white background. GO number GSE180004 and GSE180818.
